# Epstein-Barr virus promotes T cell dysregulation in a humanized mouse model of multiple sclerosis

**DOI:** 10.1101/2022.02.23.481716

**Authors:** Jessica R. Allanach, Naomi M. Fettig, Blair K. Hardman, Vina Fan, Ariel R. Rosen, Erin J. Goldberg, Zachary J. Morse, Iryna Shanina, Galina Vorobeychik, Lisa C. Osborne, Marc S. Horwitz

## Abstract

Infection with the human-tropic Epstein-Barr virus (EBV) is a strong risk factor for multiple sclerosis (MS), though the underlying mechanisms remain unclear. To investigate the immunomodulatory effects of latent EBV infection, we induced experimental autoimmune encephalomyelitis (EAE) in immunocompromised mice humanized with peripheral blood mononuclear cells (PBMCs) from individuals with or without a history of EBV infection and/or a diagnosis of relapsing MS. HuPBMC EAE mice generated from EBV seronegative healthy donors were less susceptible to developing severe clinical disease than EBV seropositive healthy donor and RRMS cohorts. Donor EBV seropositivity and RRMS led to a significant incremental increase in the number of brain and spinal cord infiltrating effector T cells, in the absence of viral reactivation, due to enhanced proliferation of donor T cells and reduced regulatory T cell expansion. The data indicate that a history of EBV infection, further compounded by a diagnosis of RRMS, promotes T cell-mediated disease in a novel humanized mouse model of MS.

**SUMMARY:** In a novel humanized mouse model of multiple sclerosis (MS), donor history of Epstein-Barr virus (EBV) infection exacerbates disease severity by skewing the balance of effector and regulatory T cells in the brain and spinal cord. These results reveal an immunomodulatory mechanism by which latent EBV infection could predispose to the development of autoimmune disease.

## INTRODUCTION

Multiple sclerosis (MS) is an autoinflammatory disease of the central nervous system (CNS) characterized by lesions of demyelinated nerve axons, infiltration of autoreactive T cells, and subsequent neurodegeneration (Dendrou, Fugger and Friese, 2015). Clinical presentation among affected individuals is heterogeneous, with variable symptoms and disease courses ranging from relapsing-remitting (RRMS) to progressive forms with intermediate phenotypes (Klineova and Lublin, 2018). Symptoms also vary widely but typically involve some combination of loss of coordination and sensation, muscle weakness, vision impairment, fatigue, pain, cognitive impairment, and others (Compston and Coles, 2008). MS is especially prevalent among women, for whom there is a heavy bias in incidence (Rommer *et al*., 2020). Although the exact etiology of MS is unclear, development of the disease is thought to be a consequence of environmental triggers in genetically susceptible individuals (Olsson, Barcellos and Alfredsson, 2017).

The environmental factor most strongly and consistently linked to MS is infection with Epstein-Barr virus (EBV) (Ascherio *et al*., 2001; DeLorenze *et al*., 2006; Thacker, Mirzaei and Ascherio, 2006; Handel *et al*., 2010; McKay *et al*., 2016; Bjornevik *et al*., 2022; Lanz *et al*., 2022). EBV is a human gammaherpesvirus transmitted through saliva that infects over 90% of all adults (Hjalgrim, Friborg and Melbye, 2007). During acute infection the virus infects epithelial cells in the oropharynx and subsequently disseminates and infects memory B cells, within which the virus establishes latency and persists for the life of the host (Hjalgrim, Friborg and Melbye, 2007; Chaganti *et al*., 2009). EBV reactivates intermittently to transmit to new cells and hosts but is typically quiescent and, in healthy individuals, does not cause disease. However, EBV is associated with a broad spectrum of malignancies and diseases (Hjalgrim, Friborg and Melbye, 2007; Chen, 2011). The varied pathological consequences of EBV infection are considered to be due to host-specific differences in immune responses to infection dictated by age, genetic predispositions, and confounding environmental exposures (Hjalgrim, Friborg and Melbye, 2007; Balfour *et al*., 2013; Houldcroft and Kellam, 2015; Sallah *et al*., 2020; Chen *et al*., 2022). Acute infection is usually asymptomatic when the virus is acquired at an early age, but delayed acquisition can cause infectious mononucleosis (IM); a self-limiting clinical condition marked by a large expansion of CD8^+^ T cells specific to viral antigens (Dunmire, Hogquist and Balfour, 2015). The connection between EBV infection and MS was first posited due to the prominent epidemiological overlap in cases of IM and MS (Warner and Carp, 1981). Numerous studies have since provided evidence linking EBV infection with the development of MS (Ascherio *et al*., 2001; DeLorenze *et al*., 2006; Thacker, Mirzaei and Ascherio, 2006; McKay *et al*., 2016; Bjornevik *et al*., 2022; Lanz *et al*., 2022).

Individuals who are EBV seronegative (EBV^-^) have a relatively low risk for MS, whereas asymptomatically infected individuals have a moderate risk, and those who have previously developed IM have a significantly higher risk for MS than those who acquired the virus asymptomatically (Thacker, Mirzaei and Ascherio, 2006; Handel *et al*., 2010). Additionally, nearly all adult and pediatric MS patients are EBV seropositive (EBV^+^) (Alotaibi *et al*., 2004; Banwell *et al*., 2007; Levin *et al*., 2010), and MS patients show higher EBV-specific antibody titers than age-matched controls in the years preceding disease onset (DeLorenze *et al*., 2006). MS patients also show increased frequencies of EBV-specific T cells in the blood compared to unaffected individuals with a similar viral load, some of which are cross reactive to myelin epitopes (Lünemann *et al*., 2006, 2008; Lanz *et al*., 2022). Recently, a longitudinal case-matched study demonstrated a 32-fold increased risk for MS and significant upregulation of a biomarker of axonal degeneration in individuals with MS, specifically following EBV seroconversion in young adulthood (Bjornevik *et al*., 2022). The efficacy of B cell depletion therapies in MS patients further suggests a role for EBV infection in disease, as anti-CD20 antibodies may target the viral reservoir (Jagessar *et al*., 2013). Moreover, EBV infection is associated with the development of other autoimmune diseases, including rheumatoid arthritis and systemic lupus erythematosus (Balandraud and Roudier, 2018; Jog and James, 2021), which suggests a common mechanism by which EBV predisposes for autoimmunity. Latent gammaherpesvirus infection is known to modulate immune responses relevant to autoimmunity, including downregulation of B cell apoptosis, suppression of regulatory T cell (Treg) function and production of an interferon-I response (Gasper-Smith, Marriott and Bost, 2006; Quan *et al*., 2010; Pratt, Zhang and Sugden, 2012). Despite mounting epidemiological and clinical evidence suggesting EBV infection is a causative factor in MS, experimental investigation of mechanism has been limited due to the high prevalence of EBV in the general population, the preponderance of asymptomatic EBV infection years prior to MS onset, and, since EBV exclusively infects humans, a lack of suitable animal models.

Due to the narrow tropism of latent EBV for human B cells, and resulting inability to infect rodent immune cells, the animal models that have helped shaped our understanding of MS pathogenesis are not amenable to EBV infection directly. To address this species limitation, our group previously developed a mouse model wherein wild-type C57Bl/6 mice are latently infected with the rodent homolog of EBV, gammaherpesvirus 68 (γ-HV68), and induced with experimental autoimmune encephalomyelitis (EAE), a widely used autoimmune model of MS, to mimic the viral acquisition observed in humans years prior to disease onset (DeLorenze *et al*., 2006; Miller, Karpus and Davidson, 2007; Olivadoti *et al*., 2007; Casiraghi *et al*., 2012, 2015; Bjornevik *et al*., 2022). Mice latently infected with γ-HV68 prior to EAE induction experience a severe pathology more alike MS than uninfected EAE mice (Casiraghi *et al*., 2012, 2015). The enhanced MS-like disease is characterized by greater clinical scores with earlier symptom onset, as well demyelinated lesion formation in the brain, and CNS infiltration of activated macrophages/microglia, cytotoxic CD8^+^ T cells, and Th1-skewed CD4^+^ T cells (Casiraghi *et al*., 2012). Treg frequencies are also significantly decreased in the CNS and spleens of infected EAE mice (Casiraghi *et al*., 2012). Moreover, these effects were exclusive to latent γ-HV68 infection, as acute γ-HV68 infection and chronic infection with other viruses, including murine cytomegalovirus and lymphocytic choriomeningitis virus, were not able to enhance EAE (Casiraghi *et al*., 2012). The immunopathological features of the γ-HV68/EAE model are hallmark characteristics of MS that are either far less pronounced or absent in classical EAE models (Simmons *et al*., 2013; Wagner *et al*., 2020), which strongly indicates EBV might contribute to similar aspects of the pathogenesis of MS in humans. Although this modeling approach has been educational, a more complete understanding of how EBV modifies MS susceptibility could be gained from improved models of host-viral interactions.

Here, we sought to study the role of EBV latency in MS using humanized mice that contain reconstituted human immune systems naturally exposed to EBV. EBV infection of humanized mice has recapitulated various aspects of the viral immune response and pathogenesis of EBV-associated diseases observed in humans, including lytic and latent infections (Melkus *et al*., 2006; Yajima *et al*., 2008; Lin *et al*., 2015), lymphoproliferation and tumor formation (Islas-Ohlmayer *et al*., 2004; Lim *et al*., 2007; Heuts *et al*., 2014), among others (Kuwana *et al*., 2011; Sato *et al*., 2011; Münz, 2015). Studies of EBV infection in humanized mice, however, have not reported CNS-localized inflammation leading to motor deficits, as seen in classical EAE models of MS. To capture the effects of long-term latent EBV infection on immune function and neuroinflammation, we induced EAE in immunodeficient mice reconstituted with peripheral blood mononuclear cells (PBMCs) isolated from healthy EBV^+^ or EBV^-^ adults. Since the vast majority of people living with MS are EBV^+^ (Levin *et al*., 2010; Abrahamyan *et al*., 2020), we also tested donor PBMCs from this group for neuroinflammatory potential in EAE. Supporting a role for EBV-mediated immunomodulation in MS susceptibility, we found that donor EBV serostatus clearly defined the time of EAE symptom onset and severity of disease. Further, this novel mouse model revealed that PBMCs from EBV seropositive donors led to diminished regulatory T cell expansion and enhanced effector T cell infiltration of the CNS, providing new mechanistic understanding of how latent EBV infection promotes CNS autoimmunity.

## RESULTS

### EBV seropositive and RRMS donor derived PBMCs exacerbate clinical disease severity in HuPBMC EAE mice

To evaluate the impact of EBV infection on clinical outcomes of disease in a humanized MS model, we engrafted NOD/SCID-IL2Rγ ^-/-^ (NSG) mice with PBMCs (HuPBMC mice) from three donor groups based on EBV seropositivity and RRMS diagnosis (Figure 1A). For this study, we enrolled treatment naïve female donors aged 19–39, due to the increased prevalence of RRMS among young women (Rommer *et al*., 2020). Consistent with previous reports (DeLorenze *et al*., 2006; Mouhieddine *et al*., 2015; Bjornevik *et al*., 2022), the donors diagnosed with RRMS were all EBV seropositive (EBV^+^) and exhibited significantly elevated EBV-specific serum IgG titres to both the acute phase viral capsid antigen (VCA, Figure 1B) and the latent phase Epstein-Barr nuclear antigen 1 (EBNA-1, Figure 1C) compared to previously infected, otherwise healthy EBV^+^ donors. Uninfected EBV seronegative (EBV^-^) healthy donors (HDs) were identified and grouped based on the absence of serum IgM and IgG specific to both antigens. EBV viral load analysis of donor PBMCs confirmed that none of the donors had detectable levels of cell-associated EBV that might be indicative of a highly active infection or an EBV-associated disorder such as malignancy or infectious mononucleosis (Kimura *et al*., 1999) (Figure 1D). We also assessed donor group differences in other serological factors associated with MS (McKay *et al*., 2016; Olsson, Barcellos and Alfredsson, 2017). Our donors showed 33–50% seropositivity for cytomegalovirus (CMV) within each donor group, which reflects the general population (Staras *et al*., 2006; Lachmann *et al*., 2018) (Table 1). Although there was no significant difference in anti-CMV IgG titres between EBV^+^ HD and the other two groups, RRMS EBV^+^ donors had elevated anti-CMV IgG compared to EBV^-^ HD (Figure S1A). We also verified that the donors did not have any existing overt seroreactivity to the inducing antigen, recombinant human myelin oligodendrocyte glycoprotein (rhMOG, Figure S1B), and the donor groups had statistically similar serum 25-OH vitamin D levels (Figure S1C). Human leukocyte antigen (HLA) typing revealed that two of four RRMS donors expressed the MS risk allele DRB1*15:01 (Wang *et al*., 2020), whereas none of the healthy donors expressed this allele (Figure S1D). Individual donor characteristics, including clinical MS parameters and viral serostatus, are reported in Table 1.

**Figure 1.**
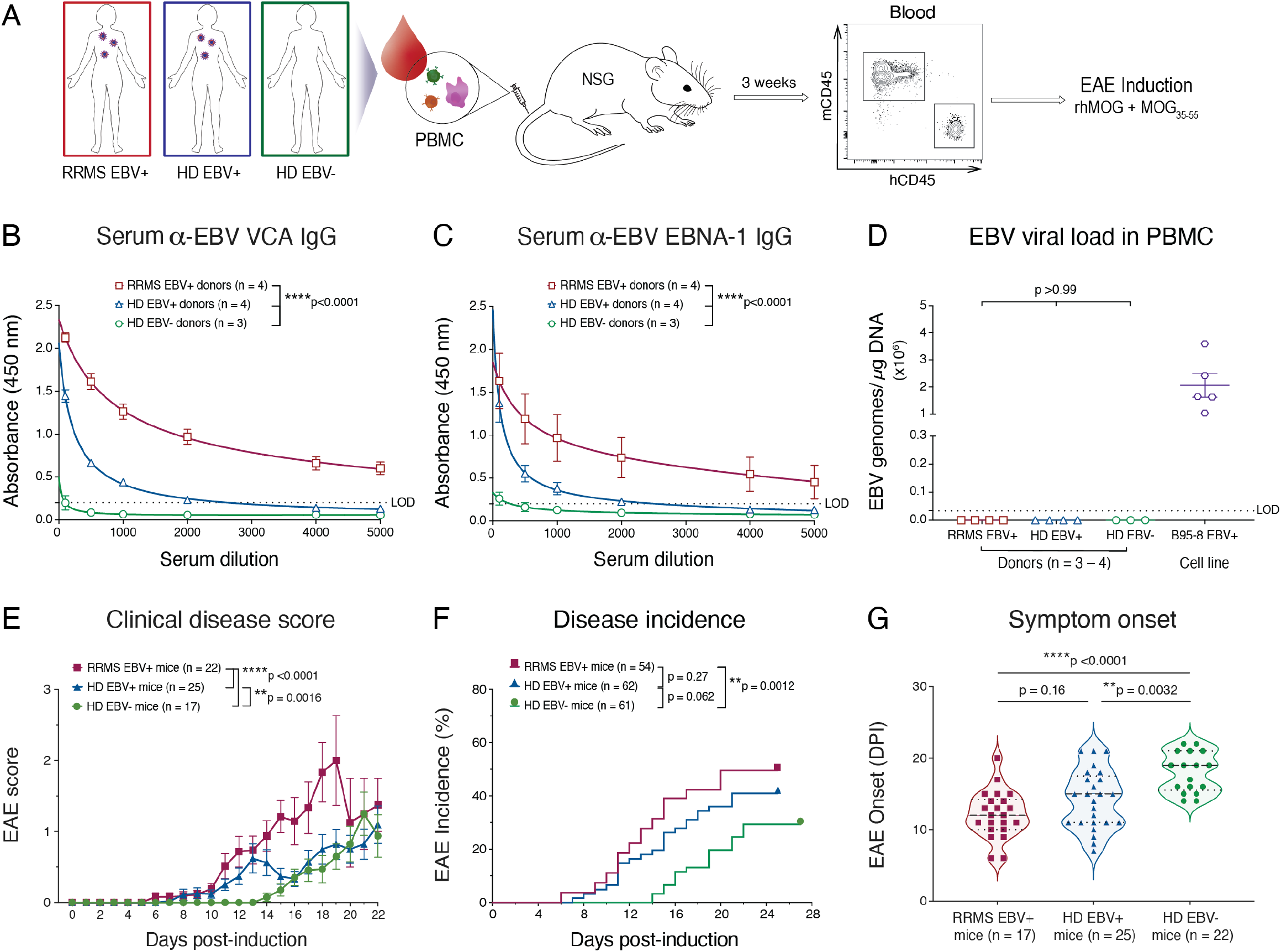
Donor EBV infection and RRMS diagnosis worsen clinical outcomes in HuPBMC EAE mice. (A) Experimental design: Donor PBMCs isolated from women with or without a history of EBV infection or an RRMS diagnosis were used to engraft immunocompromised NSG mice. Following a three-week reconstitution period and confirmation of circulating human CD45^+^ cell repopulation, humanized NSG mice (HuPBMC) were immunized with recombinant human myelin oligodendrocyte glycoprotein (rhMOG) antigens to induce EAE. (B) Donor serum IgG specific to acute EBV antigen viral capsid antigen (VCA). (C) Donor serum IgG specific to latent EBV antigen Epstein-Barr nuclear antigen 1 (EBNA-1). For B and C, group data are shown as mean with SEM and were curve fit with a one-site total binding equation. Statistical differences in titre curves were assessed by ordinary two-way ANOVA. (D) Cell-associated EBV viral loads in donor PBMCs. Data are shown as mean with SEM and were analyzed by ordinary one-way ANOVA with Tukey’s multiple comparisons test. For B – D, n = 3 – 4 donors/group and the lower limit of detection (LOD) for each assay is represented by a dashed line. (E) Clinical disease scores post-induction for symptomatic HuPBMC EAE mice. Data are shown as mean with SEM and curves were analyzed by ordinary two-way ANOVA (n = 17 – 25 mice/group derived from 3 – 4 donors/group). (F) Incidence of clinical EAE symptoms post-induction. Data are shown as percentage of the group and curves were analyzed by Log-rank (Mantel-Cox) test (n = 54 – 62 mice/group derived from 3 – 4 donors/group). (G) Day of EAE symptom onset post-induction (DPI). Distribution of individual data are shown with median and quartiles (dashed lines) and were analyzed by Brown-Forsythe and Welch ANOVA with Dunnett’s T3 multiple comparisons test (n = 17 – 25 mice/group derived from 3 – 4 donors/group).

**Table 1.** PBMC donor demographics, disease characteristics, and serology. All RRMS and healthy donors were female. RRMS participants were treatment naïve and in clinical remission at the time of donation. Abbreviations: Healthy donor, HD; Expanded Disability Status Scale, EDSS; Viral capsid antigen, VCA; Epstein-Barr nuclear antigen 1, EBNA-1; Cytomegalovirus, CMV.

| Donor ID | Age (years) | Age (years) at MS symptom onset | EDSS score | Disease duration | EBV IgG serostatus |  | CMV IgG serostatus | Serum 25-OH (ng/mL) |
| --- | --- | --- | --- | --- | --- | --- | --- | --- |
| MS-01 | 24 | 24 | 2.5 | 4 months | VCA | + | + | 29.7 |
|  |  |  |  |  | EBNA-1 | + |  |  |
| MS-02 | 31 | 31 | 2.5 | 6 months | VCA | + | + | 34.9 |
|  |  |  |  |  | EBNA-1 | - |  |  |
| MS-03 | 32 | 26 | 2.0 | 5 years | VCA | + | - | 19.9 |
|  |  |  |  |  | EBNA-1 | + |  |  |
| MS-04 | 39 | 31 | 2.0 | 7 years | VCA | + | - | 66.5 |
|  |  |  |  |  | EBNA-1 | + |  |  |
| HD-01 | 39 | N/A | N/A | N/A | VCA | + | - | 27.7 |
|  |  |  |  |  | EBNA-1 | + |  |  |
| HD-02 | 25 | N/A | N/A | N/A | VCA | - | - | 35.0 |
|  |  |  |  |  | EBNA-1 | - |  |  |
| HD-03 | 26 | N/A | N/A | N/A | VCA | - | - | 29.8 |
|  |  |  |  |  | EBNA-1 | - |  |  |
| HD-04 | 23 | N/A | N/A | N/A | VCA | + | + | 37.5 |
|  |  |  |  |  | EBNA-1 | + |  |  |
| HD-05 | 19 | N/A | N/A | N/A | VCA | - | + | 18.9 |
|  |  |  |  |  | EBNA-1 | - |  |  |
| HD-06 | 25 | N/A | N/A | N/A | VCA | + | + | 38.4 |
|  |  |  |  |  | EBNA-1 | - |  |  |
| HD-07 | 28 | N/A | N/A | N/A | VCA | + | - | 27.5 |
|  |  |  |  |  | EBNA-1 | + |  |  |

Flow cytometric analysis of transplanted PBMCs confirmed that recipient NSG mice were engrafted with a similar composition of immune cells from each donor (Figure S2). HuPBMC mice were then induced with EAE using a mixture of rhMOG_1-120_ and MOG_35-55_ after a three-week reconstitution period (Figure 1A). HuPBMC mice were found to be susceptible to EAE and exhibited symptoms typical of other murine EAE models, including weight loss, piloerection, motor imbalance, and an ascending paralysis with resolution within approximately a week from onset (data not shown and Figure S3, S7). The development of paralysis, reaching EAE scores of up to 4 on a standard 5-point scale (Miller, Karpus and Davidson, 2007), was similar to the asynchronous, relapsing EAE phenotype of the Non-Obese Diabetic (NOD) background strain for the NSG (Figure S3, S7) (Baker *et al*., 2019). HuPBMC EAE mice demonstrated an exacerbation of clinical disease by multiple measures when engrafted with PBMCs from donors who were EBV^+^ and had been diagnosed with RRMS (Figure 1E – G). The overall disease course of EAE was significantly worsened in EBV^+^ HD mice compared to EBV^-^ HD mice, which was in turn significantly more severe in the EBV^+^ RRMS recipient cohorts compared to both HD groups (Figure 1E). The incidence of EAE symptoms was significantly greater in EBV^+^ RRMS recipient mice compared to the HD groups, and moderately increased with EBV infection between HD groups (Figure 1F). HuPBMC EAE mice derived from EBV^-^ HD donors also exhibited a significantly delayed time to symptom onset compared to the EBV^+^ recipient groups, which was not significantly different between the EBV^+^ recipient groups, regardless of an RRMS diagnosis (Figure 1G). The clinical outcomes of EAE indicate that donor EBV infection and RRMS diagnosis both increase disease susceptibility and severity in recipient HuPBMC EAE mice.

### Human T cells infiltrate and localize to areas of inflammatory demyelination in the CNS of HuPBMC EAE mice

In classical EAE models, clinical disease scores are assessed based on the severity of ascending paralysis, which is a consequence of inflammatory damage to the spinal cord (Simmons *et al*., 2013). To investigate the underlying cause of the clinical symptom differences associated with donor EBV and RRMS status, we first evaluated the degree of demyelination of the spinal cord among the three recipient HuPBMC EAE groups. EAE induction of immunocompetent NOD mice and RRMS/HD EBV^+^ HuPBMC mice led to a significant loss of total myelin content relative to unaffected NSG control mice (Figure 2A). Both EBV^+^ groups had significantly less myelin in the spinal cord compared to NSG controls, whereas EAE induction of EBV^-^ HD recipient HuPBMC mice did not significantly reduce myelination, suggesting greater protection from EAE consistent with diminished clinical disease severity in this group. As demyelination in EAE is predominantly T cell driven (Furtado *et al*., 2008; Hohlfeld and Steinman, 2017), we confirmed that engrafted human T cells could infiltrate the CNS following induction. Notably, we detected an accumulation of human CD8^+^ T cells in the white matter of both the spinal cord (Figure 2B) and the cerebellum (Figure 2C) of HuPBMC EAE mice engrafted with RRMS donor PBMCs, demonstrating that engrafted human T cells can localize to CNS regions typically demyelinated in classical EAE models. Moreover, we observed an inflammatory lesion within the cerebellum marked by microgliosis, as indicated by concentrated Iba-1 staining, that co-localized with human CD8^+^ T cell infiltration (Figure 2D). Demyelination of the spinal cord and brain lesion infiltrates composed of macrophages and CD8^+^ T cells are consistent with the CNS immunopathology seen in both EAE models and MS, respectively (Salou *et al*., 2015; Wagner *et al*., 2020).

**Figure 2.**
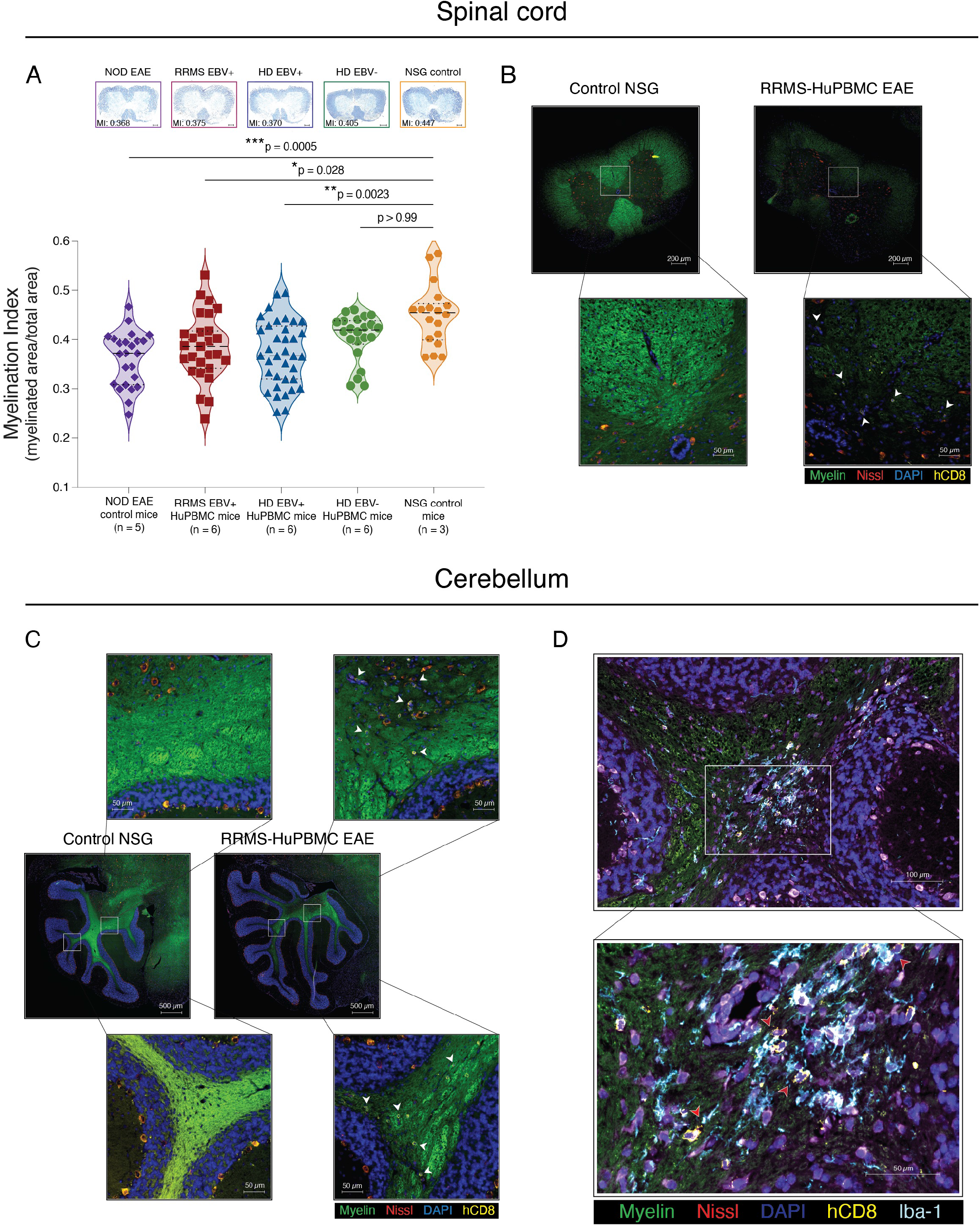
EAE induction in HuPBMC mice causes human T cell infiltration and inflammatory demyelination of the CNS. (A) Demyelination of the spinal cord in the HuPBMC EAE model. Perfused spinal cords were obtained days 19 – 25 post-induction (5 – 8 days post-symptom onset) from HuPBMC EAE mice, and days 15 – 25 post-induction from NOD EAE mice (5 – 15 days post-symptom onset). Eriochrome cyanine-stained sections (top) obtained from the lower thoracic region of the spinal cord show representative myelination indices (MI) for each of the respective group means. Individual data points represent averages of serial sections sampled from 4 – 6 equidistant regions along the entire length of the spinal cord (n = 18 regional points from 3 unengrafted NSG control mice; n = 22 – 36 regional points from 5 – 6 mice/group for EAE-induced NOD and HuPBMC groups). Distribution of individual data are shown with median and quartiles (dashed lines) and were analyzed by Kruskal-Wallis with Dunn’s multiple comparisons test. (B – D) Human CD8^+^ cells infiltrate the CNS of HuPBMC EAE mice engrafted with RRMS donor PBMCs. Representative images of lumbar spinal cord (B) and cerebellar (C, D) sections from an unengrafted control NSG mouse (left) and a symptomatic HuPBMC EAE mouse (right) derived from a donor with RRMS. Perfused tissues were collected day 15 post-EAE induction (day 4 post-symptom onset). Sections were labelled with FluoroMyelin (green), NeuroTrace 530/615 (red), DAPI (blue), anti-hCD8 (yellow) and anti-Iba-1 (light blue). Example hCD8^+^ cells are indicated by white arrows (B, C), and hCD8^+^ cells in proximity to Iba-1^+^ cells by red arrows (D). Scale bars indicate size as specified per panel, showing (A) 200 μm; (B) 200 μm, insets 50 μm; (C) 500 μm, insets 50 μm; and (D) 100 μm, inset 50 μm.

### CNS immunopathology in HuPBMC EAE is mediated by human Th1 and cytotoxic T cells and murine macrophages

Murine EAE models can present with a range of distinct disease symptomologies, CNS pathologies and constituent autoimmune processes based on the induction method and rodent strain used. As this HuPBMC EAE model is novel, we first characterized the immunopathological features of EAE mice using NSG mice engrafted with HD EBV^+^ PBMCs, in order to establish a baseline from which to compare EBV^-^ and RRMS PBMCs recipient cohorts. Consistent with our findings by histology, we measured substantial spinal cord infiltration of both human CD4^+^ (hCD4^+^, 3802 ± 3525 cells/ spinal cord) and CD8^+^ (hCD8^+^, 1225 ± 1262 cells/ spinal cord) T cells by flow cytometry in HuPBMC EAE mice (Figure 3A). A large proportion of hCD4^+^ T cells in the CNS produced IFNγ (∼45%). Although there were relatively few IL-17A^+^ cells, the majority of these cells co-expressed IFNγ, indicating that this model is predominantly Th1-driven (Figure 3A, B). Among CNS-infiltrating hCD8^+^ T cells, ∼80% expressed IFNγ and/or Granzyme B (GzmB) (Figure 3A, B), suggesting they are capable of cytotoxic activity. Though the level of T cell reconstitution and infiltration was variable among tissues derived from the same recipient cohort, the pattern of cytokine expression was similar in the periphery and in the CNS (Figure 3B). Human T cell infiltration was detected in the CNS of all EAE-induced HuPBMC mice regardless of the development of clinically visible EAE symptoms (Figure S4). However, the spinal cords of mice that developed clinical symptoms contained significantly greater numbers of hCD45^+^ cells, including total hCD3^+^ and activated hCD8^+^ T cells, than mice that remained subclinical (Figure S4). Moreover, there was a significantly higher ratio of hCD8^+^ T cells relative to hCD4^+^ T cells within the CNS of mice with EAE symptoms (Figure 3C), suggesting a critical role for cytotoxic hCD8^+^ T cells in mediating demyelination and tissue damage.

**Figure 3.**
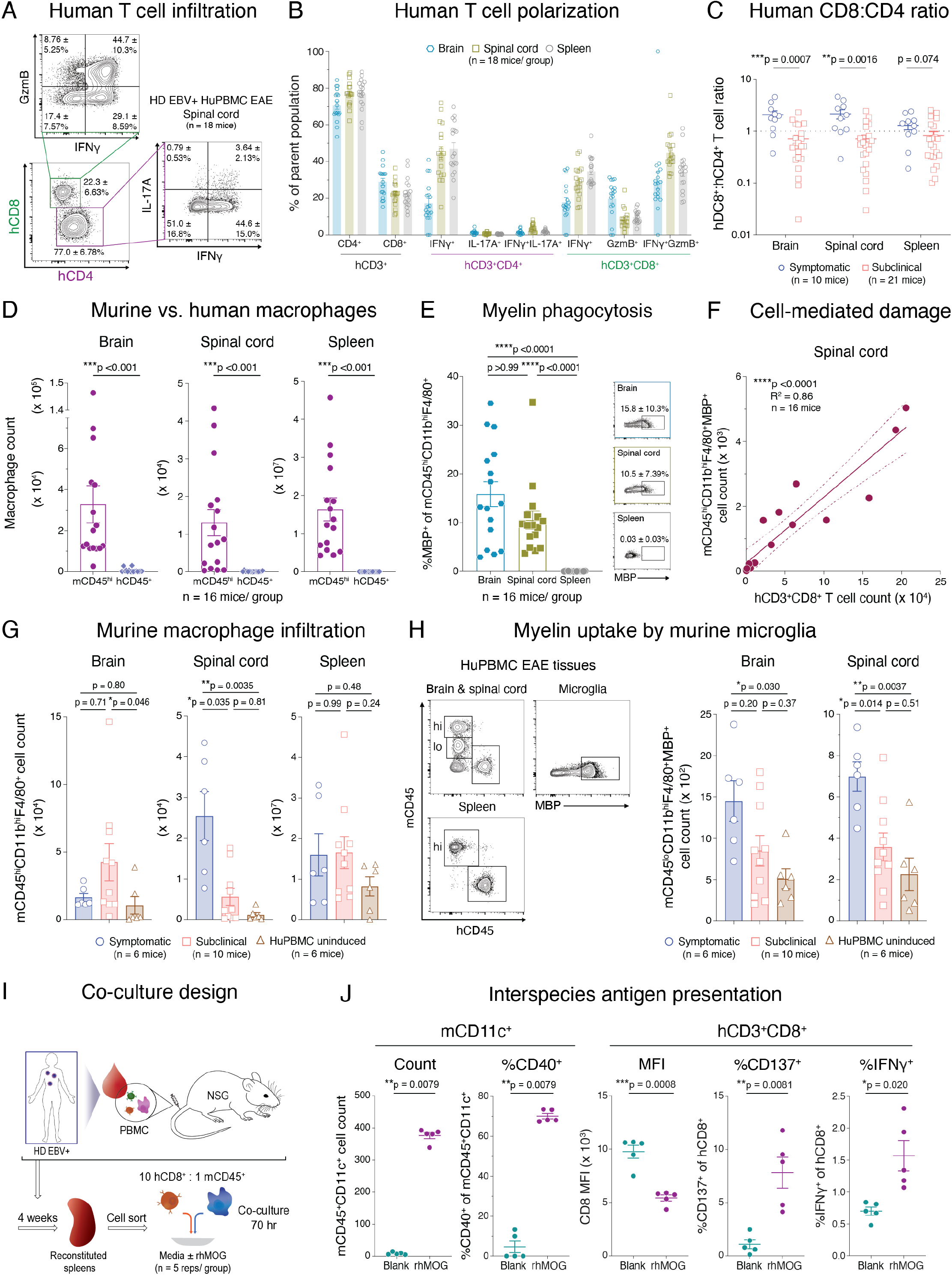
EAE in HuPBMC mice is mediated by infiltrating human Th1 and cytotoxic T cells and by murine myeloid cells. (A) Concatenated flow cytometric plots of spinal cord-infiltrating hCD3^+^CD4^+^ and hCD3^+^CD8^+^ T cells and their respective cytokine expression, shown as mean frequency of the parent population ± SD. (B) Human T cell subsets quantified in the brain, spinal cord and spleen, shown as mean with SEM. For A and B, perfused tissues were collected day 22 post-EAE induction and isolated cells were stimulated with PMA and ionomycin (n = 18 mice derived from one EBV^+^ HD). For C – H, perfused tissues were collected days 14 – 24 post-induction of recipient cohorts derived from 2 – 3 unrelated EBV^+^ HD PBMCs, and data were combined for analysis. (C) Human CD8^+^:CD4^+^ T cell ratios in the CNS and spleens of HuPBMC EAE mice that either developed symptoms or remained subclinical. Data are shown as mean with SEM (n = 10 – 21 mice/group) and were analyzed by Mann-Whitney test. (D) Total numbers of murine mCD45^hi^CD11b^hi^F4/80^+^ and human hCD45^+^CD14^+^CD68^+^ macrophages in the CNS and spleens of HuPBMC EAE mice. Data are shown as mean with SEM (n = 16 mice/group) and were analyzed by Mann-Whitney test. (E) Frequency of murine macrophages containing intracellular myelin basic protein (MBP) in HuPBMC EAE organs. Data are shown as mean with SEM (n = 16 mice/group) and were analyzed by Kruskal-Wallis with Dunn’s multiple comparisons test. Concatenated flow cytometric plots, right, show the mean frequency of the parent population ± SD. (F) Correlation between the total numbers of infiltrating hCD3^+^CD8^+^ T cells and murine macrophages containing intracellular MBP in the spinal cords of HuPBMC EAE mice. Data were analyzed by simple linear regression (n = 16 mice). Goodness of fit is indicated by R^2^ value, and the 95% confidence interval by dashed lines. (G) Total numbers of infiltrating mCD45^hi^CD11b^hi^F4/80^+^ macrophages in the CNS and spleens of uninduced HuPBMC mice and EAE-induced HuPBMC mice that either developed symptoms or remained subclinical. Data are shown as mean with SEM (n = 6 – 10 mice/group). CNS groups were analyzed by Kruskal-Wallis with Dunn’s multiple comparisons test and spleen groups by Brown-Forsythe and Welch ANOVA with Dunnett’s T3 multiple comparisons test. (H) Total numbers of murine microglia (mCD45^lo^CD11b^hi^F4/80^+^) containing intracellular MBP (gating scheme left) in the CNS of uninduced HuPBMC mice and EAE-induced HuPBMC mice that either developed symptoms or remained subclinical. Data are shown as mean with SEM (n = 6 – 10 mice/group) and were analyzed by Brown-Forsythe and Welch ANOVA with Dunnett’s T3 multiple comparisons test. (I) Interspecies co-culture assay design: NSG mice (n = 4) were engrafted with PBMC from an EBV^+^ HD, and after four weeks, reconstituted spleens were processed and combined for fluorescence-activated cell sorting (FACS). Sorted cells were cultured 1 mCD45^+^:10 hCD8^+^ cells per well in blank or rhMOG-supplemented complete culture medium for 70 hours. (J) Changes in mCD11c^+^ and hCD8^+^ cell marker expression following myelin antigen exposure in culture. Data are shown as mean with SEM (n = 5 replicate wells/group) and analyzed by Mann-Whitney test (mCD11c^+^ data) or by unpaired t-test with Welch’s correction (hCD8^+^ data). Median fluorescence intensity (MFI).

Myelin phagocytosis by infiltrating and resident macrophages/microglia is a key mediator of demyelination in EAE models (Yamasaki *et al*., 2014; Dong and Yong, 2019). In humanized mouse models, both endogenous murine and engrafted human myeloid cells reconstitute and can co-exist within the CNS (Gorantla, Gendelman and Poluektova, 2012; Zhang *et al*., 2021). We assessed the abundance of murine (mCD45^hi^CD11b^hi^F4/80^+^) and human (hCD45^+^CD14^+^CD68^+^) macrophages in the CNS and periphery to determine the potential relative contribution of each subset to the immunopathology of HuPBMC EAE. Unlike hematopoietic stem cell engraftment-based humanized mouse models, PBMC humanization of NSG mice generally favors the engraftment of T cells over innate immune subsets (Morillon *et al*., 2020). Accordingly, we observed numerous mCD45^hi^ macrophages in the brains, spinal cords, and spleens of HuPBMC EAE mice but very few hCD45^+^ macrophages (Figure 3D). The overall scarcity of hCD45^+^CD14^+^CD68^+^ cells in the CNS suggests that infiltrating murine macrophages and resident microglia likely perform the bulk of axonal demyelination following EAE induction. We confirmed that murine myeloid cells phagocytosed myelin in the HuPBMC EAE model based on the detection of intracellular myelin basic protein (MBP) in mCD45^hi^ macrophages in the CNS after induction (Figure 3E). In the spinal cord, the total number of infiltrating hCD8^+^ T cells correlated strongly and specifically with the number of infiltrating MBP^+^ murine macrophages in the same tissue (Figure 3F), similar to MS brain lesions wherein infiltrating cytotoxic T cells and macrophages are positively correlated with the extent of axonal damage (Bitsch *et al*., 2000). An association with myelin phagocytosis was not observed for hCD4^+^ T cells or human macrophages (Figure S5). Furthermore, we observed a significant increase in the total numbers of murine macrophages and myelin-phagocytosing microglia (mCD45^lo^CD11b^hi^F4/80^+^MBP^+^) in the spinal cords of HuPBMC mice that developed clinical EAE symptoms compared to mice that remained subclinical (Figure 3G, H), supporting the notion that murine myeloid cells demyelinate the CNS sufficiently to produce clinical symptoms in this model.

Collectively, the data indicate that infiltrating hCD4^+^ T cells direct a Th1 polarized response in the CNS of HuPBMC EAE mice that leads to hCD8^+^ T cell-mediated myelin damage and subsequent phagocytosis by murine macrophages and microglia. The capability of hCD8^+^ T cells to exert myelin-directed cytotoxicity, however, was unclear in an interspecies model of immune-mediated demyelination. Based on previous work, which has demonstrated productive human T cell receptor (TCR) interactions with mouse major histocompatibility complex (MHC) in humanized NSG mice (Halkias *et al*., 2015), we assessed whether hCD8^+^ T cells could effectively interact with myelin antigen presented on mouse MHC I-expressing cells ex vivo (Figure 3I). Co-culture of sort-purified splenic mCD45^+^ and hCD8^+^ cells from HuPBMC mice in the presence of rhMOG resulted in a significant expansion of mCD11c^+^ cells, including those expressing CD40, and a simultaneous decrease in CD8 median fluorescence intensity on human CD3^+^CD8^+^ T cells (Figure 3J), indicative of increased antigen recognition (Kroger and Alexander-Miller, 2007). There was also a significant increase in the proportion of hCD8^+^ T cells expressing the antigen activation marker CD137, as well as IFNγ (Figure 3J), which affirms the functional capacity of hCD8^+^ T cells to recognize and respond to antigen in humanized mouse tissues. These data highlight the interspecies nature of the immunopathology of the HuPBMC EAE model and supports its use as a model of MS.

### Donor EBV seropositivity and RRMS diagnosis promote effector T cell accumulation in the CNS of HuPBMC EAE mice

Since the HuPBMC EAE recipient groups showed an EBV and RRMS associated increase in disease severity and demyelination, we sought to assess human immune cell infiltration of the CNS and reconstitution in the periphery to identify contributing immune subsets. In the brain, spinal cord, and spleen, HuPBMC EAE recipient cohorts followed a stepwise, increasing trend in total human CD45^+^ immune cell counts with donor EBV seropositivity and RRMS diagnosis (Figure 4A). In all HuPBMC EAE mice, relatively few human B cells infiltrated the CNS relative to T cells: approximately 25–100 hCD19^+^ B cells versus 10^3^–10^5^ hCD3^+^ T cells per spinal cord (Figure 4B, C). Total B cell infiltration of the CNS and peripheral reconstitution also did not differ between the three recipient groups (Figure 4B). EBV intermittently reactivates from latently infected memory B cells to transmit to new cells (Hadinoto *et al*., 2008), which has been suggested to potentially contribute to relapses in MS (Bar-Or *et al*., 2020). Though engrafted B cells reconstituted the spleen (approximately 10^3^ – 10^5^ hCD19^+^ B cells), EBV viral genomes were undetectable in the spleens of most mice at EAE endpoint, regardless of recipient group (Figure 4D), indicating peripheral viral reactivation from engrafted B cells is unlikely to have influenced disease outcomes. We further assessed whether an active EBV infection may play a role in promoting disease by measuring antibody production by EBV-specific B cells. Given that engrafted lymphocytes do not form organized germinal centers in NSG mouse lymphoid tissues, and are subsequently unable to promote class-switching and hypermutation of human B cells (Jangalwe *et al*., 2016; Lang *et al*., 2017), any EBV-specific IgG produced in these mice would indicate a reactivation of engrafted memory B cells rather than a novel response to infection post-engraftment. We confirmed the production of rhMOG-specific human IgM and the absence of rhMOG-specific human IgG in the serum of HuPBMC EAE mice (Figure S6A), indicating the engraftment of otherwise functionally responsive human B cells. In three separate EBV^+^ recipient cohorts, no human IgG to the EBV antigen VCA could be detected in serum after EAE induction (Figure S6B), despite a strong response in corresponding donor serum (Figure 1B), suggesting that, under these conditions, an active infection likely does not play a role in the enhancement of clinical disease.

**Figure 4.**
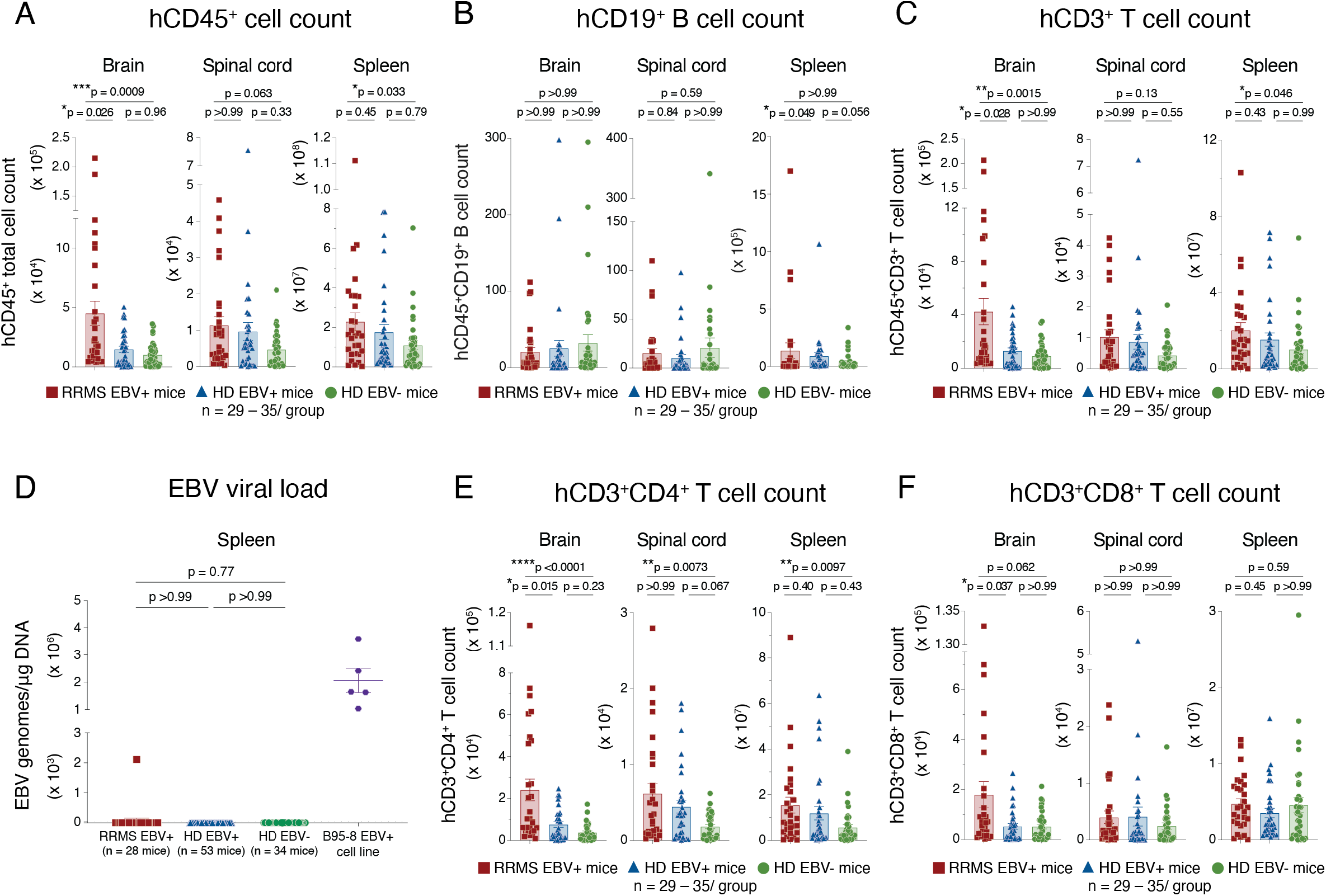
Donor EBV and RRMS status promote human immune cell infiltration of the HuPBMC EAE CNS without viral reactivation. (A) Total human CD45^+^ immune cell counts, (B) hCD19^+^ B cell counts, (C) hCD3^+^ T cell counts, (D) EBV genome copies in splenic DNA, (E) hCD3^+^CD4^+^ T cell counts and (F) hCD3^+^CD8^+^ T cell counts in the brain, spinal cord, and spleen of recipient HuPBMC EAE mice at endpoint, grouped by PBMC donor EBV serostatus and RRMS diagnosis. Perfused organs were collected days 14 – 27 post-EAE induction (average 5 – 10 days post-symptom onset). For total immune cell quantification, n = 29 – 35 mice/group derived from 2 – 3 donors/group. For viral load quantification, n = 28 – 53 mice/group derived from 2 – 4 blood donors/group; n = 5 replicates for control EBV^+^ B95-8 cell line and assay lower limit of detection is represented by a dashed line. All data are shown as mean with SEM and were analyzed by Kruskal-Wallis with Dunn’s multiple comparisons test.

Though direct B cell involvement in causing disease differences seemed improbable, total hCD3^+^ T cell counts comprised the majority of infiltrating human CD45^+^ cells in the CNS, following the same increasing trend of higher counts with donor EBV and RRMS status (Figure 4C). Among infiltrating T cells, hCD4^+^ T cells demonstrated a consistent increase in total counts in all three tissues with donor EBV seropositivity and RRMS (Figure 4E), while hCD8^+^ T cell counts were increased only in the brain of RRMS recipient mice compared to HD mice (Figure 4F). Increased hCD8^+^ T cell counts in the RRMS recipient brains corresponded to greater numbers of infiltrating murine immune cell in these samples (Figure S6C), whereas microglia and human macrophage counts did not differ between recipient groups in any tissue (Figure S6D, E). Splenic reconstitution of hCD8^+^ T cells was also similar between groups (Figure 4F).

Among hCD4^+^ T cells, the proportions of cells expressing cytokines indicative of subset skewing (Th1 and Th17) did not follow any consistent trends with PBMC donor EBV or RRMS status (Figure 5A, S7A – C), indicating polarization toward a particular inflammatory Th subset was not the main driver of clinical group differences. However, the increased number of total hCD4^+^ T cells in EBV^+^ HD and RRMS recipient group tissues than EBV^-^ HD tissues (see Figure 4E) led to an overall greater abundance of effector Th1 cells in the CNS and spleen (Figure 5B), despite similar levels of IFNγ expression (Figure 5A, S7A). The relatively fewer total numbers of Th17 (Figure 5C) and IFNγ^+^IL-17γ^+^Th cells (Figure 5D) showed some moderate, but not striking group specific trends that would otherwise suggest they underpinned the observed clinical disease differences. Despite similar numbers of total infiltrating hCD8^+^ T cells in the CNS between recipient cohorts (see Figure 4F), donor EBV seropositivity led to significantly enhanced T cell cytotoxicity (Figure 5E), as measured by a significant shift from less IFNγ expression (Figure 5F) to more GzmB expression (Figure 5G, H) among hCD8^+^ T cells in the brain and spinal cord. Interestingly, this trend was not further enhanced by donor RRMS diagnosis. Although total effector hCD8^+^ T cell counts were similar between recipient group tissues (Figure S7D – F), the skewing toward increased cytotoxic capacity within the infiltrating hCD8^+^ T cell population is consistent with the elevated level of demyelination in the CNS of EBV^+^ recipient mice (see Figure 2A). Overall, both a donor history of EBV infection and a diagnosis of RRMS led to incremental increases in the accumulation of pathogenic effector hCD4^+^ and hCD8^+^ T cells in the CNS of HuPBMC mice induced with EAE.

**Figure 5.**
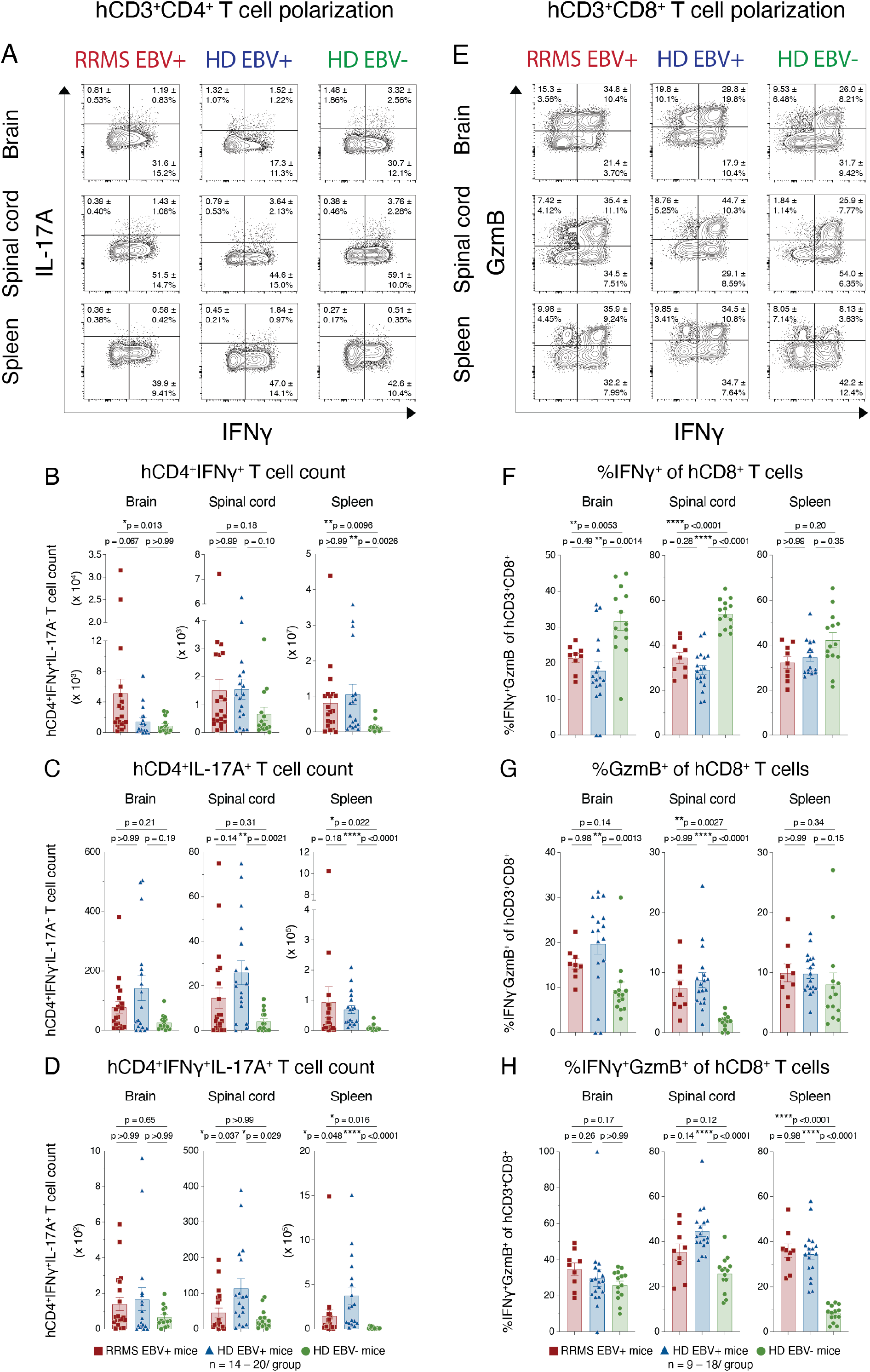
Donor EBV and RRMS status promote effector T cell expansion in the HuPBMC EAE model. Figure shows brain and spinal cord infiltration and spleen reconstitution in recipient HuPBMC EAE mice at endpoint, grouped by PBMC donor EBV serostatus and RRMS diagnosis. (A) Concatenated flow cytometric plots of IFNγ and IL-17A expression, showing the mean frequency of hCD3^+^CD4^+^ cells ± SD, as well as corresponding total (B) IFNγ^+^(IL-17A^-^), (C) IL-17A^+^(IFNγ^-^), and (D) IFNγ^+^IL-17A^+^ hCD4^+^ T cell counts in each tissue. (E) Concatenated flow cytometric plots of IFNγ and GzmB expression, showing the mean frequency of hCD3^+^CD8^+^ cells ± SD, quantified as (F) %IFNγ^+^(GzmB^-^), (G) %GzmB^+^(IFNγ^-^), and (H) %IFNγ^+^GzmB^+^ of hCD3^+^CD8^+^ cells in each tissue. Perfused organs were collected days 14 – 27 post-EAE induction (average 5 – 10 days post-symptom onset). Isolated immune cells were stimulated with PMA and ionomycin for cytokine detection (n = 9 – 20 mice/group derived from 1 – 2 donors/group). All plotted data are shown as mean with SEM and were analyzed by Brown-Forsythe and Welch ANOVA with Dunnett’s T3 multiple comparisons test or by Kruskal-Wallis with Dunn’s multiple comparisons test.

### Donor EBV seropositivity and RRMS diagnosis limit regulatory T cell expansion in HuPBMC EAE mice

Alongside greater Th1 and cytotoxic T effector cell infiltration of the CNS, we observed a reduction in the proportion of hCD4^+^ T cells expressing the regulatory transcription factor FOXP3 in EBV^+^ recipient group tissues compared to the EBV^-^ HD group (Figure 6). Initially, FOXP3 expression in hCD4^+^ T cells from freshly isolated PBMCs is comparable between donor groups (Figure 6A) but is incrementally reduced in EBV^+^ HD and RRMS hCD4^+^ T cells in the blood of NSG mice following engraftment of the PBMCs (Figure 6B). Following EAE induction, this trend of lower proportions of FOXP3^+^ T cells is likewise observed in the CNS and spleen of EBV^+^ HD and RRMS recipient mice (Figure 6C – E), even though total regulatory T cell (Treg) counts are similar between groups (Figure S6F). The sum consequence of this incremental increase in pathogenic T cell infiltration of the CNS and simultaneous suppression of Treg expansion was a significantly greater ratio of effector hCD4^+^ and hCD8^+^ T cells to regulatory hCD4^+^ T cells in the CNS and periphery of EBV^+^ HD and RRMS recipient mice compared to EBV^-^ HD mice (Figure 6F, G & S6G, H). As sample collection was timed to maintain similar overall cumulative EAE scores between groups at endpoint, differences in CNS infiltration could not be correlated with greater clinical disease burden for RRMS and EBV^+^ recipient groups (Figure S8A – D). Though weight loss followed the EBV and RRMS status trend with clinical disease severity (Figure S8E), the incidence of clinical symptoms of graft-versus-host disease (GvHD) was comparably low between the groups (Figure S8F, G), suggesting excessive, nonspecific inflammation due to varying degrees of human PBMC recognition of murine MHC within the recipient tissues was not the main determinant of the observed Th1 expansion and Treg suppression. The data collectively suggest that a history of both EBV infection and an RRMS diagnosis among donors leads to a compounded reduction in the expansion of Tregs and a concomitant increase in effector Th1 and cytotoxic T cell abundance, and, subsequently, to systemic dysregulation of T cell-mediated inflammation resulting in excessive CNS damage and worsened clinical outcomes.

**Figure 6.**
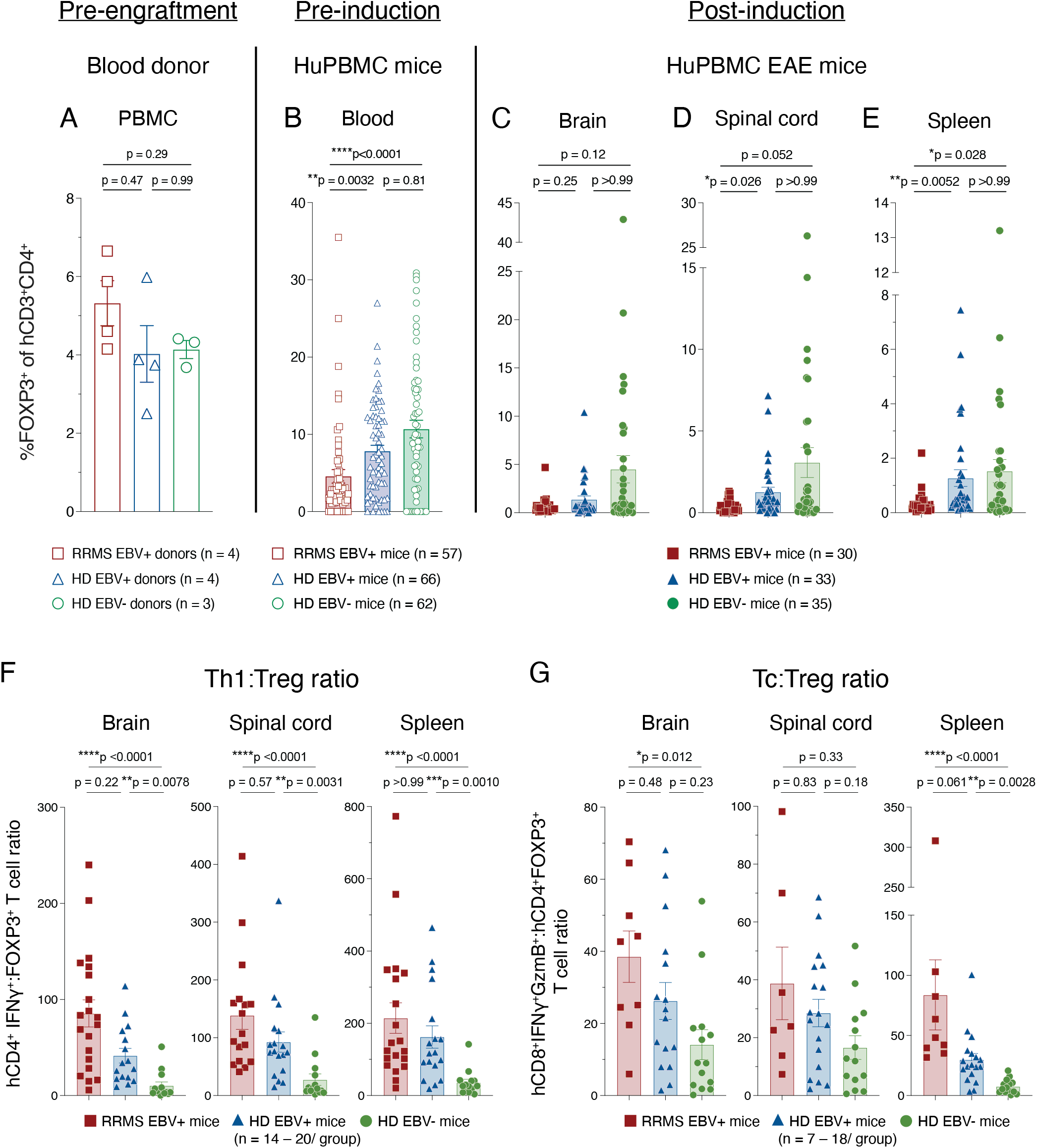
Donor EBV and RRMS status limit regulatory T cell expansion in HuPBMC EAE mice. The proportion of hCD3^+^CD4^+^ T cells expressing FOXP3 in (A) freshly isolated donor PBMC (n = 3 – 4 donors/group), (B) in the peripheral blood of engrafted HuPBMC mice at 3 weeks post-PBMC injection (n = 57 – 62 mice/group derived from 3 – 4 donors/group), and in the (C) brain, (D) spinal cord, and (E) spleen of HuPBMC EAE mice at endpoint (n = 30 – 35 mice/group derived from 2 – 3 donors/group). The ratio of infiltrating (F) hCD4^+^IFNγ^+^ (Th1) and (G) hCD8^+^IFNγ^+^GzmB^+^ (Tc) to regulatory hCD4^+^FOXP3^+^ (Treg) cells per tissue in recipient HuPBMC EAE mice at endpoint, grouped by PBMC donor EBV serostatus and RRMS diagnosis (n = 7 – 20 mice/group from 1 – 2 donors/group). Cells were isolated from perfused organs collected days 14 – 27 post-EAE induction (average 5 – 10 days post-symptom onset) and, for cytokine detection, stimulated with PMA and ionomycin. Data are shown as mean with SEM and were analyzed by Brown-Forsythe and Welch ANOVA with Dunnett’s T3 multiple comparisons test (A) or by Kruskal-Wallis with Dunn’s multiple comparisons test (B – G).

### EBV infection and RRMS both enhance donor T cell proliferation following TCR stimulation

Based on the findings above, PBMCs from EBV^-^ HD are seemingly less reactive to myelin antigen challenge in the HuPBMC EAE model than EBV^+^ HD and RRMS PBMCs. The observation that FOXP3 expression among hCD4^+^ T cells is reduced even prior to EAE induction suggests the effector T cells may become more activated during reconstitution due to a lack of regulation, which then transfers to the CNS after MOG immunization. Within the donor PBMCs (prior to engraftment into NSG mice), hCD4^+^ and CD8^+^ T cells did not differ in their group baseline expression of the activation markers HLA-DR, CD38, CD137, or CD154 (Figure S9), nor of subset-specific transcription factors (Figure S10). Likewise, baseline cytokine expression did not follow any EBV or RRMS associated trends that would explain the observations in the HuPBMC EAE mice (Figure S10). To evaluate the possibility that T cells from an EBV-exposed host are intensely activated following TCR engagement generally, we stimulated donor PBMCs by anti-hCD3/CD28 stimulation in vitro (Figure 7). Interestingly, direct stimulation of the TCR without a specific antigen resulted in a trend of enhanced hCD4^+^ T cell proliferation with donor EBV and RRMS status, as measured by a CFSE-based proliferation index and Ki-67 expression (Figure 7A – C). hCD4^+^ T cells from EBV^+^ HD PBMCs, and even more so EBV^+^ RRMS PBMCs, were able to attain a higher average number of cellular divisions during the incubation period compared to EBV^-^ donor PBMCs (Figure 7B). Consistent with the HuPBMC EAE model data, these T cells were not differentially polarized after stimulation based on inflammatory cytokine expression (Figure 7D, E). Moreover, the same effects were observed with hCD8^+^ T cells, as measured by an incremental increase in proliferation with donor EBV seropositivity and RRMS diagnosis following anti-CD3/CD28 treatment (Figure 7F – H). Stimulated hCD8^+^ T cell cytokine expression also did not follow group specific trends (Figure 7I, J). Therefore, in addition to an EBV and RRMS associated deficiency in the expansion of regulatory T cells in the HuPBMC EAE model, both hCD4^+^ and hCD8^+^ T cells from EBV experienced hosts are also more proliferative following general TCR stimulation, without alteration to the polarization profiles of these cells.

**Figure 7.**
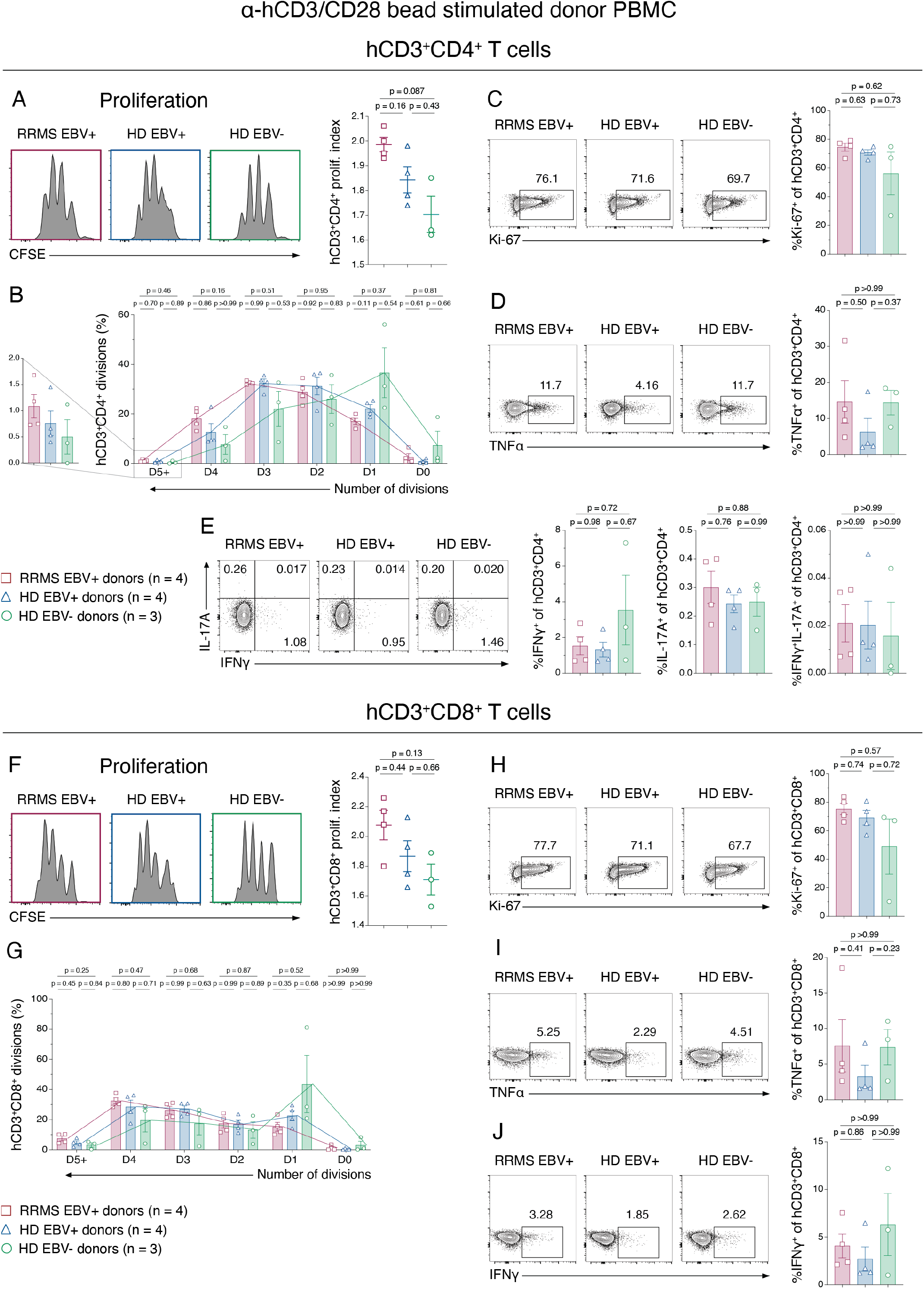
Donor T cell proliferation is enhanced by both EBV seropositivity and RRMS. Previously frozen, whole PBMC samples from EBV^+^ RRMS, EBV^+^ HD, and EBV^-^ HD blood donors were incubated with anti-CD3/CD28 coated beads for 96 hours to stimulate T cells in the absence of specific antigen. Figure shows (A) the proliferation index determined by CFSE staining, (B) the proportion of hCD4^+^ T cells having undergone a specified number of cellular divisions by CFSE staining, (C) Ki-67 expression, (D) TNFα expression and (E) IFNγ and IL-17A expression on hCD3^+^CD4^+^ T cells, as well as, (F) the proliferation index determined by CFSE staining, (G) the proportion of hCD8^+^ T cells having undergone a specified number of cellular divisions by CFSE staining, (H) Ki-67 expression, (I) TNFα expression and (J) IFNγ expression on hCD3^+^CD8^+^ T cells. Concatenated flow plots indicate the sum proportion of marker positive cells for all donors in each group. The colored symbol legend is applicable to all comparisons (n = 3 – 4 blood donors/group). All plotted data are shown as mean with SEM and were analyzed by Brown-Forsythe and Welch ANOVA with Dunnett’s T3 multiple comparisons test or by Kruskal-Wallis with Dunn’s multiple comparisons test.

## DISCUSSION

The findings of this study highlight the utility of PBMC-humanized EAE mice in modelling clinical and immunopathological aspects of MS and its association with a history of EBV infection. Immunocompromised NSG mice engrafted with human donor PBMCs were susceptible to the development of EAE symptoms and demyelination, much like classical murine EAE models immunized with myelin antigens. As confirmed by histological analysis, engrafted human CD8^+^ T cells were able to migrate to and infiltrate the mouse CNS and localize to areas of microgliosis and myelin damage. Moreover, the CNS of symptomatic mice was enriched for hCD8^+^ relative to hCD4^+^ T cells, which is commonly observed in post-mortem analysis of MS lesions (Babbe *et al*., 2000; Salou *et al*., 2015) but is not usually observed in classical murine EAE models (Simmons *et al*., 2013). We also observed equivalent numbers of infiltrating human T cells in the brain and the spinal cord, as opposed to the spinal cord dominant pathology of most murine EAE models (Simmons *et al*., 2013). Infiltrating T cells produced predominantly Th1 and cytotoxic molecules that functioned alongside murine macrophages to demyelinate the CNS. Notably, demyelination of the mouse CNS occurred without substantial involvement of engrafted human macrophages, and human CD8^+^ T cells and murine immune cells were able to functionally interact ex vivo. Interspecies TCR:MHC interactions are potentially less efficient than autologous antigen presentation (Halkias *et al*., 2015), as evidenced by a relatively reduced rate of EAE symptom incidence compared to classical murine EAE models (Dang *et al*., 2015). These interactions are, however, sufficient to produce demyelinating disease resulting in clinically measurable motor deficits. Though a previous study incorporated myelin antigen-pulsed autologous dendritic cells (DC) to provide human MHC for presentation in PBMC humanized mice (Zayoud *et al*., 2013), our data suggest human DC priming is not necessary to produce clinical EAE symptoms in HuPBMC mice. Immune system reconstitution of NSG mice with human PBMCs does not precisely replicate the composition of the donor PBMCs, nor all the nuanced contextual and site-specific complexities of the mammalian immune system (Morillon *et al*., 2020). Nevertheless, these data collectively demonstrate that the HuPBMC EAE model is an interspecies system that exemplifies key elements of MS, and as such is a viable mouse model capable of providing novel clues for causation, biomarkers, and treatment of disease.

PBMC donor history of EBV infection and a diagnosis of RRMS both exacerbated clinical disease susceptibility and severity in HuPBMC EAE mice. Consistent with the findings of Bjornevik *et al*., 2022, clinical EAE differences could not be attributed to CMV infection status, nor to reduced serum vitamin D levels. Moreover, clinical differences associated with donor EBV seropositivity in the HuPBMC EAE model are consistent with findings that elevated humoral responses to EBV at disease onset are associated with long-term greater disease severity and higher relapse rate in individuals with MS (Comabella *et al*., 2022). Though EBV infection alone was not necessary to enable EAE development in HuPBMC mice, the EBV^+^ HD recipient group presented with an intermediate phenotype that resembled RRMS recipient mice in some ways more than the EBV^-^ HD group. Interestingly, active infection by EBV was not required to exacerbate disease severity, as demonstrated by the absence of viral replication and memory B cell responses to EBV in mice derived from donors with existing latent EBV infection. Our group has previously demonstrated that viral reactivation does not occur following EAE induction in C57Bl/6 mice latently infected with the murine EBV homolog, gammaherpesvirus-68 (γHV68), and that γHV68 does not actively infect the CNS during EAE (Casiraghi *et al*., 2012). B cell accumulation in the CNS of HuPBMC EAE mice was also not significantly different among mice derived from donors with or without EBV and RRMS. Though relatively few B cells engraft and infiltrate the CNS compared to T cells, the possibility remains that engrafted B cells of varying specificities and functionalities, which can act both protectively and pathogenically in classical EAE models (Rojas *et al*., 2019; Jain and Yong, 2021), may also contribute to the enhancement of disease and the increased CNS inflammation observed in EBV^+^ and RRMS recipient mice. Clinical post-mortem evidence has suggested that EBV could contribute to the development and/or progression of MS through infection of CNS-infiltrating B cells and tertiary lymphoid structures, leading to local inflammation (Serafini *et al*., 2004, 2007; Pikor *et al*., 2016). With minimal B cell accumulation in the CNS at endpoint, the HuPBMC EAE model likely reflects earlier stages of MS, where B cell-containing lymphoid structures have not yet formed. An important limitation of PBMC humanized mice is the development of xenogeneic graft-versus-host disease, which limits the duration of experiments to ∼25 days post-immunization in our experience. Further investigation will be required to assess the presence and influence of EBV-infected and EBV-specific B cells HuPBMC EAE mice. Although there are multiple supported mechanisms by which EBV could promote an autoimmune environment through B cell-specific processes, our data suggests a role for T cell immunomodulation following latent infection.

HuPBMC EAE mice derived from EBV^+^ HD and RRMS recipient groups had increased accumulation of Th1 and cytotoxic T cells in the CNS compared to EBV^-^ HD groups. Additionally, EBV^+^ mice also had substantially reduced Treg expansion, indicating an impairment of immune regulation resulting from infection. The increased abundance of effector T cells relative to regulatory T cells consequently skews to promote a more inflammatory environment in mice derived from healthy donors with a history of EBV infection, which further compounds when donors have also been diagnosed with RRMS. Unlike in classical EAE models, where inbred mice are effectively genetically identical and highly susceptible to EAE immunization (Levine and Sowinski, 1973; Marín *et al*., 2014), the genetic variability of engrafted human immune cells from unrelated donors highlights the consistency and magnitude of the effect of EBV seropositivity on clinical and immunological disease outcomes in this model. Given that both clinical and immunological differences were observed between the healthy donor groups differing based on EBV serostatus, these effects are not entirely due to existing autoimmunity among donors. The increase in infiltrating T effector cells appeared to be due to a generalized enhanced proliferative capacity among T cells derived from EBV seropositive donors following TCR engagement. Further analysis will be required to determine if engrafted human T cells that proliferate in the periphery express more CNS homing molecules and chemokine receptors when derived from EBV^+^ donors, or, if upon infiltration of the CNS, EBV experienced T cells proliferate locally to a greater extent than those that are EBV naïve. Regulatory T cells control effector T cell proliferation and motility in the CNS following EAE induction (Koutrolos *et al*., 2014), and thus assessment of the functional capacity of the engrafted cytotoxic T cell and Treg populations would also provide insight into mechanisms by which EBV suppresses regulation of effector cells following antigenic challenge.

As T cells are required for constant immune control of EBV-infected B cells (Long, Meckiff and Taylor, 2019), latent infection could promote lasting change in predisposition to activation that may alter the balance of effector and regulatory responses toward a more ‘MS-like’ state. In the γHV68 EAE model, we also observe greater CD8^+^ T cell infiltration of the CNS, and particularly the brain, of latently infected mice, in addition to Th1-skewing, macrophage/microglia and DC activation, and reduced Treg proportions in the CNS and periphery (Casiraghi *et al*., 2012, 2015). Similar Th1-skewing and CD8^+^ T cell involvement was observed in latent γHV68-infected mice induced with rheumatoid arthritis (Mouat *et al*., 2021), suggesting a common role for gammaherpesviruses in the exacerbation of and predisposition for autoimmunity. Other susceptibility factors for MS, including a history of obesity, smoking and low vitamin D intake, have all been suggested to act in part by promoting T cell inflammation both directly and by impeding regulatory processes (Ascherio, Munger and Lünemann, 2012; Correale and Farez, 2015; Fisher *et al*., 2019; Marrodan *et al*., 2021). Ingelfinger et al. recently identified increased CD25 expression on transitional helper T cells as a defining phenotype of MS-derived PBMCs in a monozygotic twin study, which the authors posit could be preferentially responsive to an environmental immune challenge such as EBV infection (Ingelfinger *et al*., 2022). Zdimerova et al. also demonstrated a synergistic interaction between EBV infection and the MS risk allele HLA-DRB1*15:01 in stem cell-humanized mice, leading to increased T cell reactivity to myelin antigens (Zdimerova *et al*., 2020). Though two of four of the RRMS donors who provided samples to this study expressed the HLA-DRB1*15:01 risk allele, the absence of the variant among healthy donors in this small initial donor pool did not allow an analysis of the downstream effects of this variant on outcomes in the HuPBMC EAE model. A focus on the interplay of genetics and environmental exposures will enable more comprehensive investigation of MS risk factors and targeted therapies in the HuPBMC EAE model moving forward.

HuPBMC mice offer the possibility of pre-screening and selecting donors based on chosen genetic variants, environmental exposures, immunological conditions (Jodeleit *et al*., 2021; Unterweger *et al*., 2021), and, in the case of the HuPBMC EAE model, MS disease characteristics of interest, to create a personalized immune system model. The increased brain infiltration in HuPBMC EAE mice may also aid in the study of MS pathology. Though EBV induced immunomodulation of T cell responses is a possible mechanism for inciting MS, it is also possible that immunomodulation occurs alongside additional virus-specific mechanisms, such as infection and immortalization of autoreactive B cells, ineffective or abherent immune control of EBV reactivation, and structural mimicry of CNS antigens by viral proteins (Lang *et al*., 2002; Pender, 2003; Laurence and Benito-León, 2017; Lanz *et al*., 2022; Schneider-Hohendorf *et al*., 2022). An important next step is to assess clinical outcomes and T cell inflammation in the model when engrafted with PBMCs derived from donors with other autoimmune disorders, especially those also linked to EBV infection, such as systemic lupus erythematosus and rheumatoid arthritis (Balandraud and Roudier, 2018; Jog and James, 2021), to determine the generalizability of T cell immunomodulation by EBV in autoimmune susceptibility. HuPBMC EAE mice could be used for directly assessing disease modifying therapies specific to human immune targets, as well as indirectly by engrafting PBMCs derived from patients treated with therapies such as B cell monoclonal antibodies or EBV-specific T cells (Pender *et al*., 2018). Instances of promising results from biologic testing in preclinical EAE models that translated to worsened disease or serious side effects in humans could ideally also be identified in preclinical models with human immune systems before advancement to clinical trials (Schmidt *et al*., 2003; Suntharalingam *et al*., 2006; Jagessar *et al*., 2012; Kappos *et al*., 2014). HuPBMC mice could also be used to circumvent the generation of costly transgenic mouse lines for immunogenetic studies. Furthermore, the HuPBMC EAE model mechanistically reflects the epidemiological and clinical data associating EBV infection with increased risk of autoimmune disease.

In summary, our findings highlight that EBV-mediated risk for disease in a humanized mouse model of MS occurs prior to autoimmune challenge and that either the establishment or maintenance of latent infection likely modulates subsequent T cell reactivity to autoantigen years later. The data support prevention of EBV infection and establishment of latency as a strong avenue for MS preventative strategies.

## MATERIALS AND METHODS

Further information and requests for resources and reagents should be directed to and will be fulfilled by the Lead Contact, Marc Horwitz.

### Human participants

Blood donation by human participants was approved by the University of British Columbia’s Clinical Research Ethics Board and by the Fraser Health Authority, under protocol H16-02338. Individuals with RRMS were recruited at the Fraser Health MS Clinic (Burnaby, BC) under the supervision of Dr. Galina Vorobeychik. Unaffected, otherwise healthy donors were recruited at the Life Sciences Center (University of British Columbia). All donors were female, 19 – 39 years of age (mean age 31.5 ± 6.1 years for RRMS donors and 26.4 ± 6.2 years for HDs) and provided written informed consent prior to enrolment in the study, from November 2018 to August 2021. Donors with a definite RRMS diagnosis, according to Poser or 2010 McDonald criteria, and disease duration of less than 10 years, were confirmed as treatment naïve prior to donation (no previous use of any disease modifying therapies during lifetime). RRMS donors underwent a neurological exam the day of blood donation to assess Expanded Disability Status Scale (EDSS) score. Individuals with a progressive MS diagnosis or EDSS >4, that were male, pregnant, outside of the designated age range, or undergoing treatment were excluded from the present study.

### Donor sample processing

Blood samples were obtained by venipuncture and assigned an alphanumeric code to protect donor identity. Whole blood (80 mL) was processed for PBMC isolation by Lymphoprep (StemCell, #07801) gradient separation, according to manufacturer’s instructions, within an hour of collection in K_2_-EDTA coated vacutainer tubes (BD, #366643). Donor PBMCs were immediately injected into recipient mice. A subsample of freshly isolated PBMCs was retained for DNA isolation and flow cytometric analysis. A second subsample of PBMCs was stored in autologous plasma with 10% dimethyl sufoxide and placed in a Mr. Frosty™ Freezing Container (Thermo Fisher, # 5100-0001) for controlled cooling to -80°C, followed by transfer to liquid nitrogen for long-term storage. Serum was obtained by centrifugation of blood (15 min at 1300 x ɡ) collected in uncoated vacutainers (BD, #367820) and frozen at -80°C prior to analysis. Donor DNA samples were HLA genotyped by The Sequencing Center (Fort Collins, Colorado, USA) using GenDx NGSengine® software for HLA typing (v.2.27.1).

### Animals

Animal work was approved by the Animal Care Committee of the University of British Columbia, under regulation of the Canadian Council of Animal Care, under protocols A17-0266 and A17-0184. Adult male NSG (NOD.Cg-*Prkdc^scid^ Il2rg^tm1Wjl^*/SzJ, JAX #005557), NSG-SGM3 (NOD.Cg-*Prkdc^scid^Il2rg^tm1Wjl^*Tg (CMV-IL3,CSF2,KITLG)1Eav/MloySzJ, JAX #013062) and NOD (JAX #001976) mice, originally sourced from the Jackson Laboratory, began experiments at 6 – 14 weeks old. In preliminary experiments, we observed a greater incidence of graft-versus-host disease (GvHD) and reduced EAE symptom incidence in female NSG mice compared to male NSG mice engrafted with the same donor PBMCs (data not shown, DNS), and therefore male NSG mice were used as recipients for this study. Moreover, NSG-SGM3 mice, which differ from the NSG by transgenic expression of human hematopoietic cytokines (Nicolini *et al*., 2004; Jangalwe *et al*., 2016), exhibited similar incidence of EAE and GvHD symptoms as NSG mice engrafted with the same donor PBMCs (DNS) and were thus used interchangeably when randomized to recipient groups. Mice were bred in three facilities in Vancouver, British Columbia (BC Cancer Animal Resource Centre, Centre for Disease Modelling, and Modified Barrier Facility) to minimize facility-specific outcomes and housed in the same specific pathogen-free facility at the University of British Columbia for the duration of experiments (Modified Barrier Facility). Mice were housed in groups up to five animals per cage on corn cob bedding (Bed-o’Cobs, The Andersons) and fed ad libitum with PicoLab Rodent Diet 20 (#5053) standard irradiated chow with free access to autoclaved acidified water (pH 3 – 4). Housing rooms were set at 22 – 25°C with a humidity range of 50 – 70%. All NSG cages were kept on the same designated immunocompromised rack on a 14.5 – 9.5-hour light-dark cycle with sunrise at 5:30 am, sunset at 8:00 pm.

### Humanization and EAE induction

NSG mice were engrafted with 5×10^6^ donor PBMCs each (or blank phosphate buffered saline (PBS) for un-engrafted controls) by intravenous tail vein injection. Twelve to twenty mice per donor were randomly assigned to recipient groups based on PBMC yield and available stock on the day of blood donation. Following a three-week reconstitution period, circulating human CD45^+^ cell repopulation was confirmed by saphenous vein blood sampling and flow cytometric analysis. HuPBMC mice, as well as un-engrafted NSG and NOD control mice for some experiments, were actively induced with EAE on day 0 by subcutaneous injection of a 100 μL total volume containing 200 μg MOG_35-55_ peptide (GenScript), 100 μg recombinant human MOG (rhMOG_1-120_ prepared in-house) and 400 μg desiccated *M. Tuberculosis H37Ra* (DIFCO, #DF3114-338) emulsified in incomplete Freund’s adjuvant (DIFCO, #DF0639-60-6). On days 0 and 2 post-immunization, mice received an intraperitoneal injection of 200 ng pertussis toxin (List Biological Laboratories, #179A). The mice were monitored and scored daily for EAE symptoms. Clinical EAE scoring was based on the following 5-point scale: 0, no overt signs of disease; 0.5, partially limp tail; 1, limp tail; 1.5 limp tail and hind limb weakness; 2, loss of coordinated movements; 2.5, one hind limb paralyzed; 3, both hind limbs paralyzed; 3.5, hind limbs paralyzed and weakness in forelimbs; 4, forelimbs paralyzed; 5, moribund state or death by EAE. Clinical outcome assessors of HuPBMC EAE mice were blinded to the viral serostatus of the blood donors. Onset was defined by the occurrence of tail paralysis (score 0.5 or higher) for two consecutive days. Cumulative EAE scores were calculated by summing daily clinical scores from the day of onset until endpoint. Mice with motor impairments, weight loss and signs of illness were provided supportive care, including administration of subcutaneous fluids and cage heating, and provisions of wet food and nutritional gel. Humane endpoint was defined by an EAE score of 4 or greater, or a clinical health score of 4 or greater, as defined by the Animal Care Committee based on body condition, weight loss and signs of pain. Symptomatic onset of GvHD was defined by the appearance and persistence or worsening of skin dryness, redness, discoloration (jaundice), and/or hair loss. Due to intragroup variability in EAE symptom onset and ongoing GvHD and wasting following the resolution of EAE symptoms, HuPBMC EAE mice could not be maintained longer than approximately ten days post-symptom onset. Recipient cohorts were treated with the same experimental interventions for humanization and EAE induction and assigned at random to different endpoint analyses.

### Recombinant human MOG protein

The extracellular domain of recombinant human myelin oligodendrocyte protein (rhMOG_1-120_) was expressed from an *Escherichia coli* (BL21) vector (pQE-30) obtained from Drs. Christopher Linington and Nancy Ruddle (Breithaupt *et al*., 2003; Oliver, Lyon and Ruddle, 2003), and the protocol was modified from that of Dr. Jennifer Gommerman. The vector expresses a 15.8 kDa His-tagged rhMOG protein under the control of a lac operon. Following isopropyl ß-D-1-thiogalactopyranoside (IPTG) induction, the cells were lysed to release the protein product from inclusion bodies. The protein was subsequently purified using a Ni^2+^-His-bind resin column (5 mL HisTrap FF, Cytiva, #17525501) and the protein fractions were analyzed by SDS-PAGE gel electrophoresis and stained with Coomassie blue. The fractions containing sufficiently pure protein of the correct molecular weight were pooled and diluted to a protein concentration of <0.5 mg/mL in preparation for partial refolding in cellulose dialysis tubing (6 – 8 kDa molecular weight cut-off, MWCO). The refolded protein product was concentrated by centrifugation in 3K MWCO Amicon tubes (EMD Millipore, #UFC900324) until a final concentration of at least 4 mg/mL was reached. The final concentration was determined using a Nanodrop Lite Spectrophotometer (Thermo Scientific, firmware version 1.02) (MW = 15.5 kDa, e/1000 = 12.09). Protein samples were aliquoted, and flash frozen in liquid nitrogen prior to storage at -80°C. The purified and concentrated rhMOG protein was confirmed to elicit an encephalitogenic response by induction of EAE symptoms in C57Bl/6 mice.

### Tissue collection

Mice were euthanized at endpoint by isoflurane overdose. Blood was collected by cardiac puncture, followed by perfusion with 20 mL of sterile PBS for collection of CNS tissues. Brains, spinal cords, and spleens were dissected whole and kept in cold sterile PBS or in 10% neutral buffered formalin (NBF) for further processing. Spinal cords were straightened, placed onto filter paper, and submerged in NBF for histological analyses. The presence of intact emulsion was confirmed by examining the skin at the injection site. Serum was extracted from untreated blood samples by centrifugation (13 min at 8000 rpm) and stored at -80°C prior to analysis. To isolate peripheral blood cells, blood samples were treated with K2-EDTA for further processing. Tissues from animals in each recipient cohort were included for analysis regardless of the occurrence of clinically visible EAE symptoms to account for differences in incidence and severity among donor groups.

### Histological analysis

#### Immunohistochemistry

Perfused, intact organs collected for analysis of cellular infiltration by immunohistochemistry (IHC) were stored in 10% NBF for 2 hours at room temperature (RT). At 2 hours, brains were halved by sagittal division and incubated for another 2 hours along with the spinal cords (4 hours total in NBF at RT). CNS organs were washed in cold PBS for 1 hour at 4°C to remove remaining NBF, then placed in 15% sucrose solution overnight at 4 °C for cryoprotection. Organs were then placed in a 30% sucrose solution overnight at 4°C, then placed overnight at 4°C in a 25% sucrose solution containing 50% optimal cutting temperature compound (OCT, VWR Clear Frozen Section Compound, #CA95057-838). Cryoprotected organs were segmented in the case of spinal cords, and oriented in cryomolds containing OCT medium and frozen on a partially submerged aluminum block equilibrated in liquid nitrogen, then stored at -80°C for sectioning. Brain and spinal cord tissues were cut at -19°C into 12-µm sagittal and coronal sections, respectively, using a Shandon Cryotome^®^ FSE (Thermo), and placed onto Superfrost Plus microscope slides (Fisher Scientific, #12-550-15). Sections were taken at least 30-µm apart and obtained from one of either brain hemispheres, or from four equally distant points along the length of the spinal cord. Sections were rehydrated in PBS for 10 min at RT, followed by blocking in 3% mouse serum in PBS overnight at 4°C. Sections were thrice washed in PBS (5 min per wash) and incubated with rabbit anti-human CD8 primary antibody (stock diluted 1:200 – Table M1) and/or goat anti-mouse/human Iba-1 primary antibody (0.5 mg/mL stock diluted 1:300) in 3% mouse serum in PBS for 2 hours at 4°C. A rat IgG isotype primary (Abcam, #ab27478) was used to confirm the specificity of the secondary antibody binding to anti-human CD8. Sections were then thrice washed and incubated with AlexaFluor 680-conjugated donkey anti-rabbit IgG secondary antibody (2 mg/mL stock diluted 1:600) or DyLight594-conjugated mouse anti-goat IgG secondary antibody (1 mg/mL stock diluted 1:300) in 3% mouse serum in PBS for 2 hours at 4°C. Sections were thrice washed with PBS then incubated for 20 min at RT in PBS containing 0.2% Triton X-100 (PBT solution). CNS sections were incubated 20 min at RT in a PBT master mix containing FluoroMyelin^TM^ Green, NeuroTrace™ 530/615 and diamidino-2-phenylindole (DAPI) (BrainStain^TM^ Imaging Kit, Invitrogen #B34650). Sections were then thrice washed in PBT (15 min per wash) to remove excess stain. Slides were mounted with ProLong™ Diamond Antifade Mountant medium (Invitrogen, # P36961), sealed, allowed to cure for 24 hours at room temperature, and then stored at 4°C until imaged.

#### Demyelination

Perfused, intact spinal cords collected for analysis of myelination were stored in 10% NBF overnight at RT, then moved to 70% ethanol and sent to Wax-it Histology Services Inc. for paraffin embedding. To quantify myelination throughout the organ, a total of 6 – 9 consecutive 10-µm coronal sections were taken from of 4 – 6 equally distanced regions along the length of the spinal cord using a Shandon Finesse 325 microtome (Thermo Scientific). Sections were mounted on Superfrost Plus microscope slides (Fisher Scientific, #12-550-15), dried at 37°C for one hour, and then air dried overnight at RT. Sections were washed in 3 changes of xylene for 10 minutes each, then treated in 3 changes of absolute ethanol for 3 minutes each and hydrated in 95% ethanol for 5 minutes. Slides were incubated in eriochrome cyanine R (Sigma-Aldrich, #3564-18-9) staining solution (0.22% w/v ferric chloride, 0.5% v/v sulfuric acid and 0.2% w/v eriochrome cyanine R CI 43820) for 20 minutes, run under tap water for 30 seconds, and incubated in differentiating solution (5.6% w/v ferric chloride) for 5 – 10 minutes until only the white matter retained the stain. Slides were washed under running tap water for 5 minutes, then dehydrated in 3 changes of 100% ethanol for 30 seconds each (with agitation) and cleared in 3 changes of xylene for 30 seconds each (with agitation). Coverslips were mounted with VectaMount Permanent Mounting Medium (Vector Laboratories, #H-5000).

#### Imaging and analysis

Histology sections were imaged using a Zeiss Axio Observer 7 epi-fluorescent microscope outfitted with a motorized stage (x, y, z mobility) and five LED channels. Whole organ immunofluorescent sections were imaged at x20/0.65 NA air objective using an Axiocam 702 mono CMOS camera at room temperature. Isotype staining was used to set LED voltages and exposures across slides per imaging experiment. Immunofluorescent images were stitched and Z-stack (3 stacks of 1.2 µm each) fusion was performed using a wavelet transform in Zen2.6 Pro (blue edition, Zeiss). Brightfield images of EC-stained spinal cord sections were acquired at 10X/0.3 NA air objective magnification using an Axiocam 105 colour CMOS camera. Stitching and subsequent analysis of images was performed using Zen 2.6 Pro. For EC sections, single RBG images were saved as JPEG files and quantitative analysis of staining was performed using ImageJ 1.53c (NIH). Default color thresholding method was used to first select the area of stained myelin and then the total area of the spinal cord. Hue, saturation, and brightness thresholds were set based on fully myelinated NSG control sections and applied across all other sections. The Area measurement function was used to acquire the area of the selected regions in μm^2^, relative to a 200-μm scale bar. Myelination was expressed as fraction of myelin-stained area of total area of the spinal cord section. Replicate sections (6 – 9 consecutive sections per region) were averaged to generate 4 – 6 regional myelination indices per spinal cord.

### Flow cytometric analysis

#### Sample processing

Brains, spinal cords, and spleens were kept in sterile PBS on ice, then processed into single cell suspensions by passing the tissue through a 70-µm cell strainer with a syringe insert. Splenocyte suspensions were incubated in red blood cell lysis buffer (150 mM NH_4_Cl, 10 mM KHCO_3_, 0.1 mM Na_2_-EDTA) for 10 min on ice. Whole EDTA-treated blood samples were incubated in pre-warmed red blood cell lysis buffer for 15 min at RT. Brain and spinal cord samples specifically processed for the detection of intracellular MBP within phagocytic cells were predigested with 0.5 mg/mL Collagenase/Dispase^®^ (Roche, #11097113001), 0.02 mg/mL DNase I (Sigma-Aldrich, #D5025) and 2% FBS in 10 mL RPMI per sample for 45 min shaking at 180 rpm and 37°C. CNS cells were further isolated by resuspension in 40% isotonic Percoll^TM^ solution (Cytiva, #17089101) and centrifugation (15 min at 1400 rpm) to remove lipid debris. Samples analyzed for cytokine expression were incubated at 37°C (5% CO_2_) for 4 hours at 1 – 2×10^6^ splenocytes or total isolated CNS cells per 200 µL of stimulation media, containing 10% FBS, 10 ng/mL phorbol 12-myristate 13-acetate (PMA, Sigma-Aldrich, #P1585), 500 ng/mL ionomycin (Thermo Fisher, #I24222) and 1 µL/mL GolgiPlug™ Protein Transport Inhibitor (containing Brefeldin A, BD Biosciences, #555029). Cell suspensions were washed between each step with sterile PBS containing 2% FBS (FACS buffer).

#### Cell sorting and co-culture

For interspecies antigen presentation assays, splenocytes were isolated from EBV^+^ HD PBMC-engrafted male NSG mice at 4 weeks post-injection, as described above. Splenocytes were combined, stained with fluorochrome-labelled anti-mCD45 and anti-hCD8 antibodies (Table M1), and positively selected using a BD Influx^TM^ cell sorter. Sorted cells were resuspended to 4×10^4^ mCD45^+^ cells with 4×10^5^ hCD8^+^ cells (1:10 ratio) per well in 200 μL complete culture medium (RPMI-1640 plus FBS, L-glutamine, penicillin/streptomycin, MEM NEAA, sodium pyruvate, HEPES, β-mercaptoethanol), supplemented with or without 20 μg/mL rhMOG protein, and incubated for 65 hours at 37°C (5% CO_2_). Cells were then refreshed with pre-warmed culture medium (same respective rhMOG treatments) containing 1 µL/mL GolgiPlug™ Protein Transport Inhibitor and incubated for another 5 hours at 37°C (5% CO_2_). Following 70 hours total incubation time, cells were washed with serum-free PBS and stained for flow cytometric analysis as described below.

#### T cell bead stimulation assay

Donor PBMC samples previously stored in autologous plasma with 10% dimethyl sulfoxide were thawed from liquid nitrogen for activation analysis. Total PBMCs (5 x 10^6^ cells/ mL) were stained with 2.5 mM carboxyfluorescein succinimidyl ester (CFSE, Sigma #21888-25MG-F) in the dark at 4°C for 8 minutes prior to quenching with newborn calf serum. An unstimulated and PMA/ionomycin-stimulated subset of each sample was analyzed for baseline marker expression (PMA stimulation conditions described above under ‘sample processing’). Duplicate wells of 200,000 CFSE-stained cells per sample were then incubated for 91 hours at 37°C (5% CO_2_) at a 1:2 bead-cell ratio with Human T-Activator CD3/CD28 DynaBeads^TM^ (Thermo Fisher, #11161D) and 1 U/mL of recombinant human IL-2 (BioLegend, #589102) in 200 μL of complete culture medium (see above for media components). The media did not contain any factors to specifically influence polarization or activation. Cells were then refreshed with pre-warmed culture medium containing 1 µL/mL GolgiPlug™ Protein Transport Inhibitor and incubated for another 5 hours at 37°C (5% CO_2_). Following 96 hours total incubation time, cells were washed with serum-free PBS and stained for flow cytometric analysis as described below.

#### Antibody staining and analysis

Cells were stained with 1X eBioscience^TM^ Fixable Viability Dye eFluor^TM^ 506 (Thermo Fisher, #65-0866-14) for 20 min at 4°C in serum-free PBS. Cells were then washed in FACS buffer, and preincubated with human and mouse Fc block (Table M1) in FACS buffer for 10 minutes at RT. Cells were washed again and incubated with fluorochrome-labelled antibodies specific to extracellular makers (Table M1) in FACS buffer for 30 minutes at 4°C. Cells were washed and treated with transcription factor fixation and permeabilization reagent (Thermo Fisher, #00-5521-00) for 30 min at 4°C, followed by a wash with permeabilization buffer (Thermo Fisher, #00-8333-56). Cells were incubated with fluorochrome-labelled antibodies specific to intracellular makers (Table M2) in permeabilization buffer for 30 min at RT. Cells were washed with permeabilization and FACS buffer and resuspended in FACS buffer with 2 mM EDTA for acquisition. Frozen human PBMC samples were stained as detailed above for use as titration, compensation, and gating controls for all experiments. Stained cell suspensions were acquired on an Attune NxT flow cytometer (Thermo Fisher) and analyzed using FlowJo™ v10.8 Software (BD Life Sciences).

### Serology

Endogenous antibodies to EBV, CMV and rhMOG were detected using indirect enzyme-linked immunosorbent assays (ELISA). Nunc Maxisorp 96-well microtiter ELISA plates (Thermo Fisher, #439454) were coated overnight at 4°C with 1 μg/well of the peptide or protein of interest (Table M3, produced by ScenicBio) in 0.05 M carbonate buffer. For EBV EBNA-1 and CMV, epitope peptides 1 and 2 were mixed prior to well coating. The following day, the plates were washed thrice with PBS and wash buffer (PBS + 0.05% Tween-20), followed by a two-hour blocking step at RT using wash buffer + 3% bovine serum albumin. Plates were washed again, then incubated with serum samples serially diluted in blocking buffer to generate duplicate 6-point curves (1:100 – 1:5000 for human donor serum and 1:50 – 1:2500 for mouse serum). After a two-hour RT incubation with serum dilutions, the plates were washed, then incubated with HRP-labelled anti-human IgG antibody (Table M1) diluted 1:3000 in blocking buffer, or HRP-labelled anti-human IgM diluted 1:4000, for 1 hour at 37°C. The plates were then washed with PBS prior to the addition of 100 μL/well TMB substrate (BD Biosciences, #555214). Fifteen minutes after the addition of substrate, 100 μL stop solution (2 N sulfuric acid) was added to each well. The plates were read at 450 nm on a VarioSkan Plate Reader (Thermo Fisher) within 10 min of adding stop solution. Donor serum at 1:1000 dilution was used to determine seropositivity to EBV and CMV antigens. The lower limit of detection for positive IgG and IgM values was set based on the level of nonspecific background signal in negative control samples (full reaction minus antigen and/or serum). Donor serum vitamin D levels were quantified using a 25-OH Vitamin D ELISA assay test kit, according to the manufacturer’s instructions (Eagle Biosciences, #VID31-K01).

### EBV viral load

Genomic DNA was isolated from a maximum of 4×10^6^ donor PBMCs or mouse splenocytes per sample using the PureLink^TM^ Genomic DNA Minikit (Invitrogen, #K1820), according to the manufacturer’s instructions, quantified using a Nanodrop Lite Spectrophotometer (Thermo Scientific, firmware version 1.02), and stored at -80°C. A *BALF5* viral DNA polymerase gene qPCR protocol adapted from Kimura et al. was used to measure EBV load in DNA samples (Kimura *et al*., 1999). The forward and reverse primer and fluorogenic probe sequences are listed in Table M4 (produced by Integrated DNA Technologies), along with the *BALF5* gene fragment sequence used for standard curve generation. qPCR reactions comprised 300 ng genomic DNA, 0.4 μM each primer, 0.2 μM probe, and 1X QuantiNova Probe PCR master mix (Qiagen, #208256). An EBV^+^ B95-8 cell line extract (100 ng per reaction) was used as a positive control for EBV detection. Blank water was used in negative control reactions. An 8-point standard curve was generated with 1 to 10^7^ *BALF5* gBlock copies per reaction. Reactions were made up in Mx3000P 96-well plates (Agilent Technologies, #401333) and data were acquired using a CFX96 Real-Time System C1000 Thermal Cycler (Bio-Rad) and CFX manager 3.1 software for HEX detection. The cycler parameters were set according to the Qiagen QuantiNova Probe PCR protocol – 2 min at 95°C to activate DNA polymerase then 45 repeats of 5 sec at 95°C and 5 sec at 60°C. Sample copies per reaction were determined by interpolation of the standard curve using triplicate-averaged reaction threshold values (C_T_), which were then used to calculate viral load. The lower limit of detection was determined by interpolating the C_T_ value for the lowest detectable standard point per plate and was normally ∼30 EBV copies/μg DNA.

### Statistical analyses

Data were collated using Microsoft Excel, then graphed and statistically analyzed using GraphPad Prism software 9.2.0 (GraphPad Software Inc.). Figure panels were composed in Adobe Illustrator V24.3. Most graphs present group means with standard error (SEM) or standard deviation (SD) unless otherwise stated in the figure legend. No statistical methods were used to predetermine sample sizes, as donor PBMC yield limited recipient cohort size. No inclusion or exclusion criteria were used for analyses and groups include all mice from the cohort regardless of the incidence of clinical EAE symptoms, unless otherwise stated. Two normally distributed groups of data were analyzed by two-tailed, unpaired t-test with Welch’s correction. For three or more groups, normally distributed data were analyzed by ordinary one-way analysis of variance (ANOVA) with Tukey’s multiple comparisons test or Brown-Forsythe and Welch ANOVA with Dunnett’s T3 multiple comparisons test. If group data did not pass the Kolmogorov– Smirnov normality test (or Shapiro-Wilk test when group N too small to compute KS distance), a nonparametric Mann-Whitney test or Kruskal-Wallis with Dunn’s multiple comparisons test was used. A Log-rank (Mantel-Cox) test was used for incidence curve analysis, simple linear regression analysis for cell count correlations, and ordinary two-way ANOVA for group comparisons with two variables (i.e., EAE scores over time and ELISA absorbance readings with serum dilution), where the column factor p value is reported. Specific statistical tests used for each assay are noted in the figure legends, along with the number of animals or replicates per group. Significance is indicated by asterisks: ****p<0.0001, ***p<0.001, **p<0.01, and *p<0.05. The data that support the findings of this study are available from the corresponding author upon reasonable request.

## Author contributions

JRA designed and performed experiments, analyzed data, and prepared the manuscript. NMF designed and performed ex vivo cellular assays. BKH designed and performed immunofluorescent imaging analysis. VF performed serum ELISAs. ARR performed myelin staining and quantification. NMF and EJG produced rhMOG protein. EJG and ZJM facilitated donor sample collection. IS coordinated phlebotomy, mouse breeding, and aided with in vivo procedures. JRA, NMF, BKH, VF, ARR, EJG, ZJM, and IS processed tissues for cellular analysis. GV recruited and assessed donors with RRMS and provided input for the study design. LCO oversaw immunofluorescent imaging and ex vivo cellular assays and provided input for the study design. JRA and MSH conceptualized the study, designed the methodology and interpreted results for discussion. MSH oversaw the study and manuscript preparation.

## Itemized contributions

Conceptualization – JRA, MSH

Investigation – JRA, NMF, BKH, VF, ARR, ZJM, EJG, IS, GV

Methodology – JRA, NMF, BKH, LCO, GV, MSH

Visualization – JRA, NMF, BKH

Writing – original draft – JRA

Writing – review and editing – JRA, NMF, BKH, LCO, MSH

Resources – LCO, GV, MSH

Supervision – LCO, GV, MSH

Funding acquisition – GV, MSH

## Funding sources

The study was supported by grants and scholarships from the Multiple Sclerosis Society of Canada (MSSC, grant ID #3631), including the endMS personnel award program (JRA, NMF), the University of British Columbia (UBC), including the UBC 4YF award (BKH, ZJM), the Canadian Institutes of Health Research (CIHR, grant ID #168967, #178298), and the National Multiple Sclerosis Society (NMSS, grant ID #PP-1707-28236). The funders had no role in study design, data collection and interpretation, or the decision to submit the work for publication.

## Declaration of interests

The authors declare no competing financial interests.

## Acknowledgements

The authors thank the following individuals for their contributions to the study: Soren Gantt and Karen Simmons for providing the EBV *BALF5* qPCR protocol; John Priatel for providing the B95-8 EBV^+^ cell line; Jennifer Gommerman, Nancy Ruddle and Christopher Linington for providing the rhMOG-producing *E. coli* plasmid; Angela Lin, Guillermo Caballero, Natalie Strynadka, and Michael Murphy for assistance with rhMOG protein production; Tanya Kadach for organizing blood donations at the Fraser Health MS Clinic; the staff at the Modified Barrier Facility for animal husbandry; Yu Gu for ELISA protocol development; Brankica Culibrk and Philma Molina for phlebotomy; Justin Wong and Andrew Johnson for assistance with FACS; Virginie Jean-Baptiste and Isobel Mouat for assistance with animal monitoring. The authors acknowledge and are thankful for technical support from the Life Sciences Institute Cores (ubcFLOW) supported by the UBC GREx Biological Resilience Initiative. The authors express their gratitude to the blood donors for providing their time and samples to the study.

---

**Table M1.** Antibodies for extracellular targets.

| Target antigen | Clone | Species reactivity | Application | Source | Identifier |
| --- | --- | --- | --- | --- | --- |
| CD3 | OKT3 | Human | Flow cytometry | Thermo Fisher | 56-0037-42 |
| CD4 | RPA-T4 | Human | Flow cytometry | BD Biosciences | 555347 |
| CD4 | RPA-T4 | Human | Flow cytometry | Thermo Fisher | 47-0049-42 |
| CD8 | SP16 | Human | IHC-F | Thermo Fisher | MA5-14548 |
| CD8 | RPA-T8 | Human | Flow cytometry | Thermo Fisher | 11-0088-42 |
| CD8 | RPA-T8 | Human | Flow cytometry | BD Biosciences | 563821 |
| CD11b | M1/70 | Mouse | Flow cytometry | Thermo Fisher | 17-0112-82 |
| CD11c | N418 | Mouse | Flow cytometry | Thermo Fisher | 11-0114-85 |
| CD14 | 63D3 | Human | Flow cytometry | BioLegend | 367142 |
| CD14 | 61D3 | Human | Flow cytometry | Thermo Fisher | 11-0149-42 |
| CD19 | HIB19 | Human | Flow cytometry | Thermo Fisher | 47-0199-42 |
| CD20 | 2H7 | Human | Flow cytometry | BioLegend | 302328 |
| CD21 | HB5 | Human | Flow cytometry | Thermo Fisher | 12-0219-42 |
| CD25 | CD25-4E3 | Human | Flow cytometry | Thermo Fisher | 11-0257-42 |
| CD25 | BC96 | Human | Flow cytometry | Thermo Fisher | 45-0259-42 |
| CD27 | LG.3A10 | Human | Flow cytometry | BioLegend | 124216 |
| CD38 | HIT2 | Human | Flow cytometry | BioLegend | 303529 |
| CD40 | 3/23 | Mouse | Flow cytometry | Thermo Fisher | MA5-17855 |
| CD45 | HI30 | Human | Flow cytometry | Thermo Fisher | MHCD4517 |
| CD45 | HI30 | Human | Flow cytometry | BD Biosciences | 557748 |
| CD45 | 30-F11 | Mouse | Flow cytometry | Thermo Fisher | 45-0451-82 |
| CD45 | 30-F11 | Mouse | Flow cytometry | Thermo Fisher | 48-0451-82 |
| CD56 | TULY56 | Human | Flow cytometry | Thermo Fisher | 17-0566-42 |
| CD68 | Y1/82A | Human | Flow cytometry | BioLegend | 333825 |
| CD137 | 4B4-1 | Human | Flow cytometry | BioLegend | 309818 |
| CD154 (40L) | 24-31 | Human | Flow cytometry | BioLegend | 310823 |
| F4/80 | BM8 | Mouse | Flow cytometry | Thermo Fisher | 47-4801-82 |
| Fc receptor<br>(CD16/CD32) | 2.4G2 | Mouse | Flow cytometry | BD Biosciences | 553142 |
| Fc receptor | Fc1.3216 | Human | Flow cytometry | BD Biosciences | 564220 |
| HLA-DR | L243 | Human | Flow cytometry | Thermo Fisher | 17-9952-42 |
| Iba-1 | Polyclonal | Mouse | IHC-F | Thermo Fisher | PA5-18039 |
| IgG (H+L) | Polyclonal | Goat | IHC-F | Jackson ImmunoResearch | 205-585-108 |
| IgG (γ chain) | Polyclonal | Human | ELISA | Thermo Fisher | 62-8420 |
| IgG (H+L) | Polyclonal | Rabbit | IHC-F | Thermo Fisher | A10043 |
| IgG2b | m2b-25G4 | Mouse | Flow cytometry | Thermo Fisher | 11-4220-82 |
| IgM (μ chain) | Polyclonal | Human | ELISA | Thermo Fisher | A18841 |

**Table M2.** Antibodies for intracellular targets.

| Target antigen | Clone | Species reactivity | Application | Source | Identifier |
| --- | --- | --- | --- | --- | --- |
| EOMES | WD1928 | Human | Flow cytometry | Thermo Fisher | 11-4877-41 |
| FOXP3 | 236A/E7 | Human | Flow cytometry | Thermo Fisher | 17-4777-42 |
| Granzyme B | QA16A02 | Human | Flow cytometry | BioLegend | 372214 |
| IFNγ | 4S.B3 | Human | Flow cytometry | Thermo Fisher | 48-7319-42 |
| IL-17A | eBio64DEC17 | Human | Flow cytometry | Thermo Fisher | 25-7179-42 |
| Ki-67 | Ki-67 | Human | Flow cytometry | BioLegend | 350516 |
| MBP | MBP101 | Mouse | Flow cytometry | Abcam | ab62631 |
| RORγt | AFKJS-9 | Human | Flow cytometry | Thermo Fisher | 12-6988-80 |
| T-bet | 4B10 | Human | Flow cytometry | Thermo Fisher | 45-5825-82 |
| TNFα | MAb11 | Human | Flow cytometry | Thermo Fisher | 17-7349-82 |

**Table M3.**
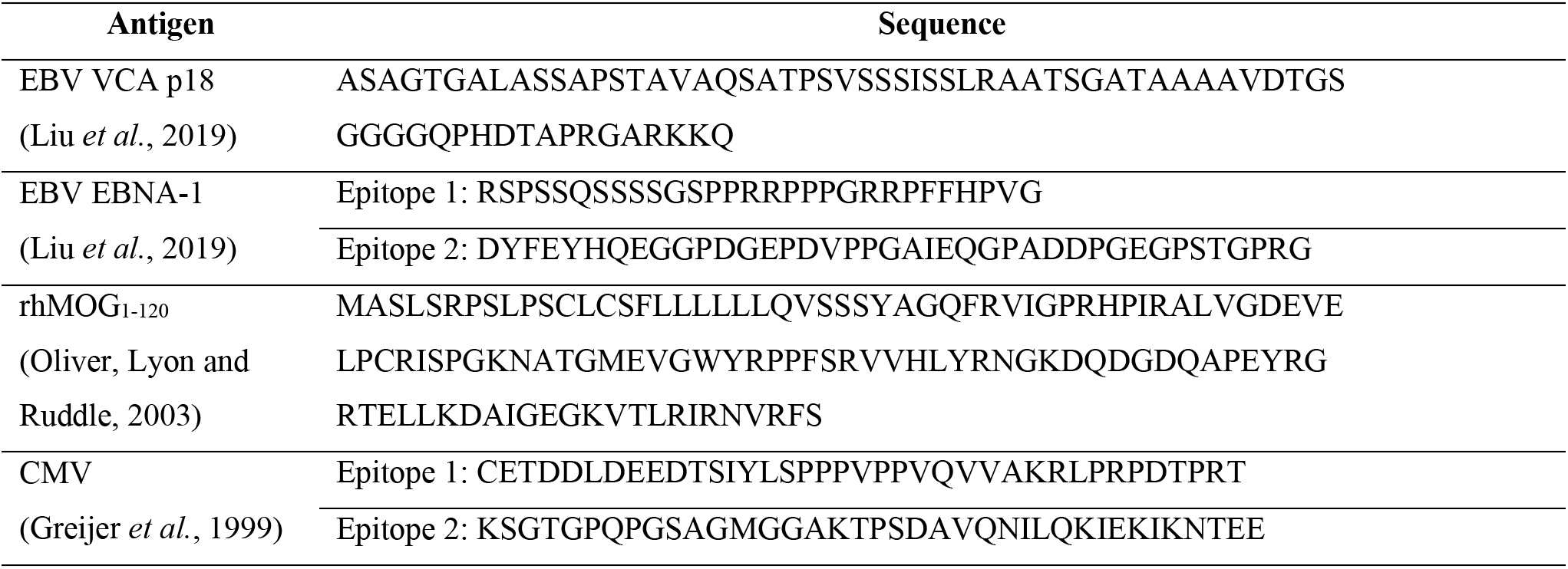
Antigen sequences for endogenous antibody detection by ELISA.

**Table M4.**
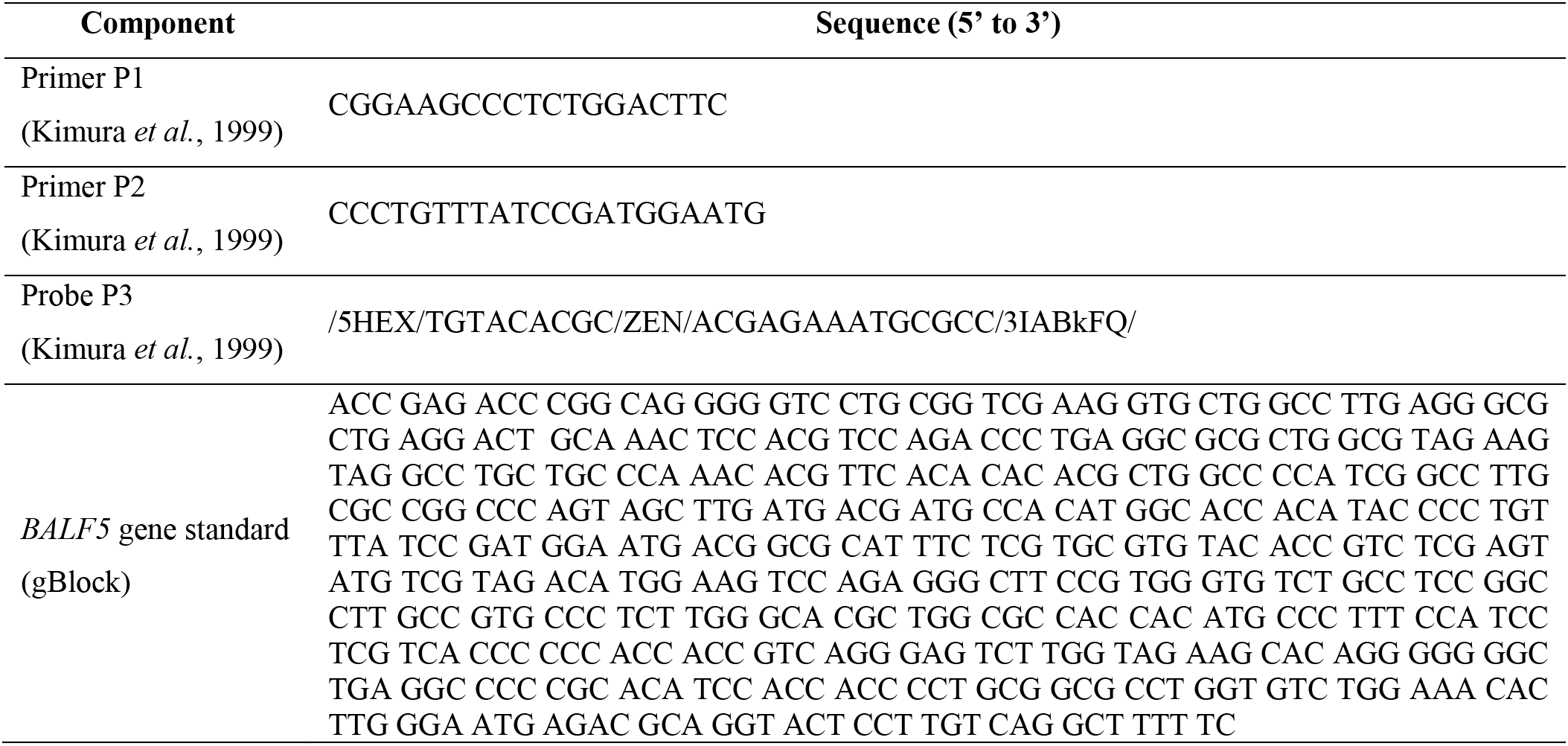
DNA sequences for EBV *BALF5* qPCR assay.

## Notes

### Competing Interest Statement

The authors have declared no competing interest.

### Summary of Updates

Updated to include additional findings

