## Supplemental information for "Epstein-Barr virus promotes T cell dysregulation in a humanized mouse model of multiple sclerosis"

**^*^ Corresponding**:

Marc S. Horwitz

Life Sciences Centre, Office 3551

2350 Health Sciences Mall

University of British Columbia

Vancouver, B. C., Canada V6T 1Z3

604-822-6298

**SUPPLEMENTAL MATERIAL**

The supplemental material included with this manuscript contains additional data cited in text relating to the main figures, pertaining to blood donor serology and immunophenotyping, HuPBMC EAE model characterization, auxiliary recipient group comparisons, and baseline donor PBMC activation and polarization.

**ABBREVIATIONS**

| CNS | Central nervous system |
| --- | --- |
| DC | Dendritic cell |
| EAE | Experimental autoimmune encephalomyelitis |
| EBNA-1 | Epstein-Barr nuclear antigen 1 |
| EDSS | Expanded disability status scale |
| GvHD | Graft-versus-host disease |
| GzmB | Granzyme B |
| HD | Healthy donor |
| HuPBMC | Human peripheral blood mononuclear cell mouse model |
| IM | Infectious mononucleosis |
| MBP | Myelin basic protein |
| (rh)MOG | (Recombinant human) Myelin oligodendrocyte glycoprotein |
| (RR)MS | (Relapsing-remitting) Multiple sclerosis |
| NOD | Non-Obese Diabetic |
| NSG | NOD/SCID-IL-2Rγc ^-/-^ (NOD.Cg-*Prkdc^scid^ Il2rg^tm1Wjl^*/SzJ) |
| PBMC | Peripheral blood mononuclear cell |
| Treg | Regulatory CD4^+^ T cell |
| VCA | Viral capsid antigen |
| γHV68 | Murine gammaherpesvirus-68 |

**SUPPLEMENTARY FIGURE LEGENDS**

**Figure S1**. **Donor serology and HLA allele sequencing for confounding MS risk factors**. Donor serum IgG specific to (A) cytomegalovirus (CMV) antigens and (B) the inducing antigen, recombinant human myelin oligodendrocyte glycoprotein (rhMOG_1-120_). For A and B, group data are shown as mean with SEM (n = 3 – 4 donors/group) and were curve fit with a one-site total binding equation. Statistical differences in titre curves were assessed by ordinary two-way ANOVA. The lower limit of detection (LOD) is represented by a dashed line. (C) Donor serum 25-OH vitamin D levels. Data are shown as mean with SEM (n = 3 – 4 donors/group) and were analyzed by Brown-Forsythe and Welch ANOVA with Dunnett’s T3 multiple comparisons test. (D) Individual human leukocyte antigen (HLA) allelic variants sequenced at three HLA-I and eight HLA-II encoding genomic loci.

**Figure S2**. **Phenotypic composition of donor PBMC injected into recipient NSG mice**. (A) The number of cells from each major human immune subset transplanted per mouse and the relative proportions of (B) T cell, (C) regulatory T cell (Treg), and (D) B cell subsets among those transplanted cells. For A – D, data are shown as mean with SEM (n = 3 – 4 donors/group) and were analyzed by Brown-Forsythe and Welch ANOVA with Dunnett’s T3 multiple comparisons test or Kruskal-Wallis with Dunn’s multiple comparisons test. Nonsignificant p values represent the overall test result for the three-group comparison, wherein each individual group comparison was also determined to be nonsignificant. The symbol legend in A is applicable to B – D.

**Figure S3**. **Clinical EAE outcomes in NOD and HuPBMC mice**. Figure shows comparative clinical outcomes following MOG_35-55_ peptide EAE immunization of wild-type Non-Obese Diabetic (NOD mice) and HuPBMC-NSG mice derived from an EBV^+^ HD. (A) Clinical disease scores over time and (B) time to EAE symptom onset days post-induction (DPI). (A, B) Data are shown as mean with SEM (n = 6 – 9 symptomatic mice/group). (C) EAE symptom incidence over time (n = 15 – 19 mice induced/group) and (D) the proportion of symptomatic mice to achieve a specific peak EAE score (n = 6 – 9 symptomatic mice/group).

**Figure S4**. **Spinal cord infiltrating human immune cell counts define clinical symptom presentation in HuPBMC EAE mice**. Total numbers of (A) hCD45^+^ immune cells, (B) hCD3^+^ T cells, (C) hCD3^+^CD8^+^ T cells, (D) activated hCD3^+^CD8^+^CD137^+^ T cells, (E) hCD3^+^CD4^+^ T cells, (F) hCD3^+^CD4^+^FOXP3^+^ regulatory T cells, (G) activated hCD3^+^CD4^+^CD137^+^ T cells, (H) hCD14^+^CD68^+^ macrophages and (I) hCD19^+^ B cells, in the CNS and spleens of uninduced HuPBMC mice and EAE-induced HuPBMC mice that either developed symptoms or remained subclinical. Perfused tissues were collected days 14 and 24 post-induction of recipient cohorts derived from two unrelated EBV^+^ HD PBMC, and data were combined for analysis. Data are shown as mean with SEM (n = 6 – 10 mice/group) and were analyzed by Brown-Forsythe and Welch ANOVA with Dunnett’s T3 multiple comparisons test or by Kruskal-Wallis with Dunn’s multiple comparisons test.

**Figure S5**. **Relationship between CNS infiltrating human T cells and murine or human macrophages in HuPBMC EAE mice**. Top row shows correlations between the total number of hCD3^+^CD8^+^ T cells and (A) murine macrophages (Mac; mCD45^hi^CD11b^hi^F4/80^+^) in the spinal cord, (B) murine macrophages in the brain containing intracellular myelin basic protein (MBP), and (C) human macrophages (hCD45^+^CD14^+^CD68^+^) in the spinal cord. Bottom row shows correlations between the total number of hCD3^+^CD4^+^ T cells and (D) murine macrophages, (E) murine macrophages containing intracellular MBP, and (F) human macrophages, in the spinal cord. Perfused tissues were collected days 14 and 24 post-induction of recipient cohorts derived from two unrelated EBV^+^ HD PBMC, and data were combined for analysis. Data were analyzed by simple linear regression (n = 16 mice). Goodness of fit is indicated by R^2^ value, and the 95% confidence interval by dashed lines.

**Figure S6**. **Human antibody production and immune cell infiltration in HuPBMC EAE mice**. (A) Deficiency in human Ig class-switching from IgM to IgG in response to the rhMOG inducing antigen in HuPBMC EAE mice (n = 8 mice derived from one HD EBV^+^ donor). (B) Absence of human IgG specific to EBV viral capsid antigen (VCA) in HuPBMC EAE mice derived from three unrelated EBV^+^ donors (n = 6 – 10 mice/group derived from one donor each). For A and B, serum samples were collected days 15 – 25 post-induction (average 5 – 8 days post-symptom onset) or from PBS-injected control NSG (n = 2 – 6 mice/group). Data are shown as mean with SEM. As most data points fell below the limit of detection (LOD, dashed line), data were not assessed statistically. (C) Murine CD45^hi^ immune cell counts, (D) murine CD45^lo^ cell counts, (E) human macrophage (hCD14^+^CD68^+^) cell counts, (F) human regulatory T cell (hCD4^+^FOXP3^+^) counts, (G) ratio of hCD8^+^IFNγ^+^ T cells to hCD4^+^FOXP3^+^ T cells, and (H) ratio of hCD8^+^GzmB^+^ T cells to hCD4^+^FOXP3^+^ T cells in the CNS and spleens of HuPBMC EAE mice. For C – H, perfused organs were collected days 14 – 27 post-EAE induction (average 5 – 10 days post-symptom onset). For total immune cell quantification, n = 29 – 35 mice/group derived from 2 – 3 donors/group. For cytokine analysis, isolated cells were stimulated with PMA and ionomycin from n = 7 – 20 mice/group derived from 1 – 2 donors/group. All individually plotted data are shown as mean with SEM and were analyzed by Brown-Forsythe and Welch ANOVA with Dunnett’s T3 multiple comparisons test or by Kruskal-Wallis with Dunn’s multiple comparisons test.

**Figure S7**. **Donor EBV and RRMS status promote effector T cell expansion in the HuPBMC EAE model, continued**. Figure shows brain and spinal cord infiltration and spleen reconstitution in recipient HuPBMC EAE mice at endpoint, grouped by PBMC donor EBV serostatus and RRMS diagnosis. (A) %IFNγ^+^(IL-17A^-^), (B) %IL-17A^+^(IFNγ^-^), and (C) %IFNγ^+^IL-17A^+^ of hCD3^+^CD4^+^ T cells, as well as, total (D) IFNγ^+^(GzmB^-^), (E) GzmB^+^(IFNγ^-^), (F) IFNγ^+^GzmB^+^ hCD8^+^ T cell counts in each tissue Perfused organs were collected days 14 – 27 post-EAE induction (average 5 – 10 days post-symptom onset). Isolated immune cells were stimulated with PMA and ionomycin for cytokine detection (n = 9 – 20 mice/group derived from 1 – 2 donors/group). All plotted data are shown as mean with SEM and were analyzed by Brown-Forsythe and Welch ANOVA with Dunnett’s T3 multiple comparisons test or by Kruskal-Wallis with Dunn’s multiple comparisons test.

**Figure S8**. **Clinical EAE and GvHD outcomes in HuPBMC EAE mice**. Figure shows (A) total average area under clinical EAE disease curves, (B) individual areas under clinical EAE disease curves, (C) distribution of cumulative EAE disease scores, (D) proportion of EAE scores attained at the peak of clinical disease, and (E) average weight loss over time, among symptomatic EAE mice (n = 17 – 25 mice/group derived from 3 – 4 donors/group), as well as the (F) incidence of symptoms of graft-versus-host disease (GvHD) over time (n = 54 – 62 mice/group derived from 3 – 4 donors/group), and (G) time to GvHD symptom onset (n = 4 – 12 symptomatic GvHD mice/group derived from 3 – 4 donors/group), among the three HuPBMC EAE recipient groups. Recipient group average EAE symptom duration to endpoint was: RRMS EBV^+^ mice 6.0 ± 1.6 days, HD EBV^+^ mice 6.9 ± 2.4 days and HD EBV^-^ mice 7.5 ± 1.4 days. For A, B, E, and G, data are shown as mean with SEM. For C, distribution of individual data is shown with median and quartiles (dashed lines). In B, C, and G, data were analyzed by Brown-Forsythe and Welch ANOVA with Dunnett’s T3 multiple comparisons test. For D, data are represented as proportions of all EAE scores attained in each group. Individual peak EAE scores per mouse were plotted and analyzed by Kruskal-Wallis with Dunn’s multiple comparisons test. In E, curves were analyzed by ordinary two-way ANOVA. For F, data are shown as percentage of the group and curves were analyzed by Log-rank (Mantel-Cox) test.

**Figure S9**. **Donor PBMC composition and baseline T cell activation**. Following donor PBMC preservation in liquid nitrogen, group differences in PBMC composition were assessed by (A) the viability of the cells (proportion alive) after recovery, (B) proportion of hCD45^+^ cells among all leukocytes, (C) proportion of hCD3^+^ T cells among hCD45^+^ cells, (D) proportion of hCD3^+^CD4^+^ T cells among hCD45^+^ cells, and (E) proportion of hCD3^+^CD8^+^ T cells among hCD45^+^ cells. Baseline donor T cell activation was assessed by the proportion of untreated hCD3^+^CD4^+^ (left) and hCD3^+^CD8^+^ (right) T cells expressing (F, K) HLA-DR (^+^, ^lo^, and ^hi^), (G, L) CD38, (H, M) both HLA-DR and CD38, (I, N) CD137, and (J, O) CD154. For F – O, concatenated flow plots indicate the sum proportion of marker positive cells for all donors in each group. All plotted data are shown as mean with SEM (n = 3 – 4 blood donors/group) and were analyzed by Brown-Forsythe and Welch ANOVA with Dunnett’s T3 multiple comparisons test or by Kruskal-Wallis with Dunn’s multiple comparisons test.

**Figure S10**. **Baseline donor T cell polarization**. Following donor PBMC preservation in liquid nitrogen, baseline donor T cell polarization was assessed by measuring the proportion of untreated (left) and PMA-Ionomycin stimulated (right) cells expressing (A) T-bet, (B) RORγt, and (C) Ki-67 on hCD3^+^CD4^+^ T cells; (D) T-bet, (E) EOMES, and (C) Ki-67 on hCD3^+^CD8^+^ T cells; (G) TNFα, (H) IFNγ, (I) IL-17A and (J) both IFNγ and IL-17A on hCD3^+^CD4^+^ T cells; and (K) TNFα and (L) IFNγ on hCD3^+^CD8^+^ T cells. Concatenated flow plots indicate the sum proportion of marker positive cells for all donors in each group. All plotted data are shown as mean with SEM (n = 3 – 4 blood donors/group) and were analyzed by Brown-Forsythe and Welch ANOVA with Dunnett’s T3 multiple comparisons test or by Kruskal-Wallis with Dunn’s multiple comparisons test.
