## Supplemental figures for "Epstein-Barr virus promotes T cell dysregulation in a humanized mouse model of multiple sclerosis"

### Figure S1

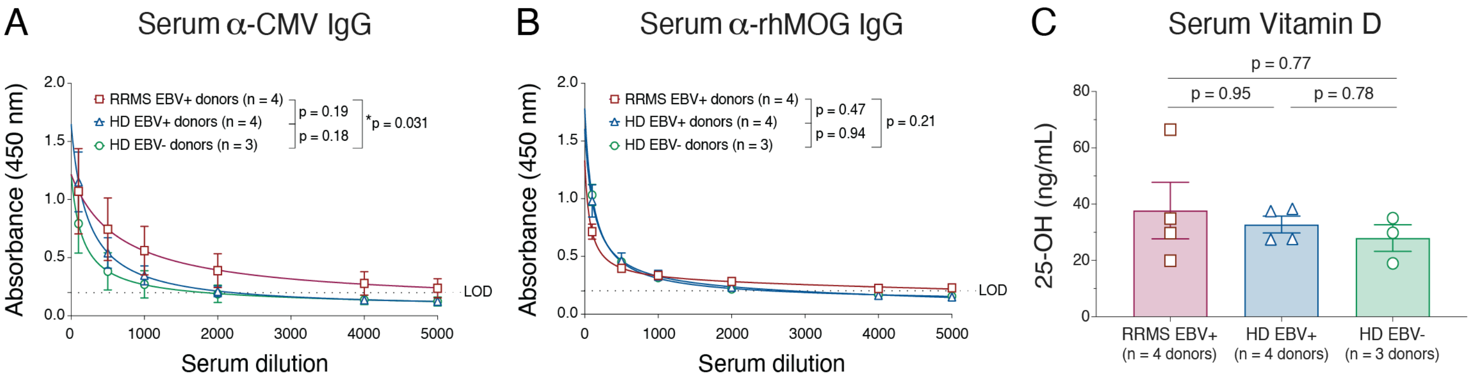


#### D Donor HLA Genotyping

| Donor ID | HLA-I | | | HLA-II | | | | | | | |
|---|---|---|---|---|---|---|---|---|---|---|---|
| | HLA-A | HLA-B | HLA-C | DRB1 | DRB3 | DRB4 | DRB5 | DQB1 | DQA1 | DPB1 | DPA1 |
| MS-01 | 02:01:01:01<br>11:01:01:01 | 41:01:01:01<br>50:01:01:01 | 06:02:01:02<br>17:01:01:05 | 03:01:01:01<br>03:01:01:01 | 02:02:01:01<br>03:01:01:01 | --<br>-- | --<br>-- | 02:01:01:01<br>02:01:01:01 | 05:01:01:03<br>05:01:01:03 | 14:01:01:01<br>104:01:01:01 | 01:03:01:02<br>02:01:01:01 |
| MS-02 | 02:01:01:01<br>02:01:01:01 | 08:01:01:02<br>08:01:01:02 | 07:02:01:01<br>07:02:01:01 | 03:01:01:01<br>03:01:01:01 | --<br>-- | --<br>-- | --<br>-- | 02:01:01:01<br>02:01:01:01 | 05:01:01:03<br>05:01:01:03 | 04:01:01:01<br>04:01:01:01 | 01:03:01:02<br>01:03:01:04 |
| MS-03 | 02:01:01:01<br>68:02:01:01 | 07:02:01:01<br>14:02:01:01 | 05:01:01:02<br>08:02:01:01 | 13:03:01:01<br>15:01:01:01 | 01:01:02:01<br>-- | --<br>-- | 01:01:01:01<br>-- | 03:01:01:03<br>06:02:01:01 | 01:02:01:01<br>05:05:01:13 | 02:01:02:01<br>04:01:01:01 | 01:03:01:01<br>01:03:01:02 |
| MS-04 | 02:01:01:18<br>26:01:01:01 | 07:02:01:01<br>35:03:01:01 | 04:01:01:01<br>07:02:01:03 | 11:15:01:01<br>15:01:17 | --<br>-- | --<br>-- | --<br>-- | 03:01:01:03<br>06:02:01:01 | 01:02:01:01<br>05:05:01:01 | 02:01:02:01<br>02:01:02:10 | 01:03:01:01<br>01:03:01:04 |
| HD-01 | 02:01:01:01<br>11:01:01:01 | 08:01:01:01<br>44:02:01:01 | 05:01:01:02<br>07:01:01:01 | 03:01:01:01<br>13:01:01:01 | 01:01:02:01<br>02:02:01:02 | --<br>-- | --<br>-- | 02:01:01:01<br>06:03:01:01 | 01:03:01:02<br>05:01:01:02 | 01:01:01:01<br>19:01:01:01 | 02:01:02:02<br>02:07:01:01 |
| HD-02 | 01:01:01:01<br>01:01:01:01 | 08:01:01:01<br>15:17:01:01 | 07:01:01:01<br>07:01:02:01 | 01:01:01:01<br>13:03:01:01 | --<br>-- | --<br>-- | --<br>-- | 03:01:01:03<br>05:01:01:03 | 01:01:01:01<br>05:05:01:13 | 03:01:01:01<br>04:02:01:02 | 01:03:01:03<br>01:03:01:05 |
| HD-03 | 01:01:01:01<br>26:01:01:01 | 08:01:01:01<br>44:03:01:01 | 07:01:01:01<br>16:01:01:01 | 03:01:01:01<br>07:01:01:01 | --<br>-- | 01:01:01:01<br>-- | --<br>-- | 02:01:01:01<br>02:02:01:01 | 02:01:01:01<br>05:01:01:02 | 04:01:01:01<br>04:01:01:01 | 01:03:01:02<br>01:03:01:02 |
| HD-04 | 01:01:01:01<br>32:01:01:01 | 08:01:01:01<br>40:02:01:01 | 02:02:02:01<br>07:01:01:01 | 03:01:01:01<br>14:01:01 | --<br>-- | --<br>-- | --<br>-- | 02:01:01:01<br>05:03:01:01 | 01:04:01:01<br>05:01:01:02 | 03:01:01:01<br>04:01:01:01 | 01:03:01:01<br>01:03:01:03 |
| HD-05 | 24:02:01:01<br>24:02:01:01 | 48:03:01:01<br>54:01:01:01 | 01:02:01:01<br>08:01:01:01 | 09:01:02:01<br>11:01:01:01 | 02:02:01:03<br>-- | 01:03:02<br>-- | --<br>-- | 03:01:01:03<br>03:03:02:02 | 03:02:01:01<br>05:05:01:01 | 05:01:01:01<br>05:01:01:01 | 01:03:01:01<br>02:02:02:01 |
| HD-06 | 01:01:01:01<br>02:01:01:01 | 08:01:01:01<br>40:01:02:01 | 03:04:01:01<br>07:01:01:01 | 07:01:01:01<br>08:03:02:01 | --<br>-- | 01:03:01:01<br>-- | --<br>-- | 02:02:01:01<br>03:01:01:17 | 02:01:01:01<br>06:01:01:03 | 04:01:01:01<br>06:01:01:01 | 01:03:01:02<br>01:03:01:03 |
| HD-07 | 02:01:01:01<br>31:01:02:01 | 27:05:02:01<br>44:02:01:01 | 02:02:02:01<br>05:01:01:02 | 12:01:01:01<br>13:02:01:02 | 02:02:01:06<br>03:01:01:01 | --<br>-- | --<br>-- | 03:01:01:05<br>06:04:01:01 | 01:02:01:04<br>05:05:01:03 | 02:01:02:01<br>03:01:01:01 | 01:03:01:01<br>01:03:01:03 |

Each cell contains alleles 1 and 2, if applicable

Figure S2

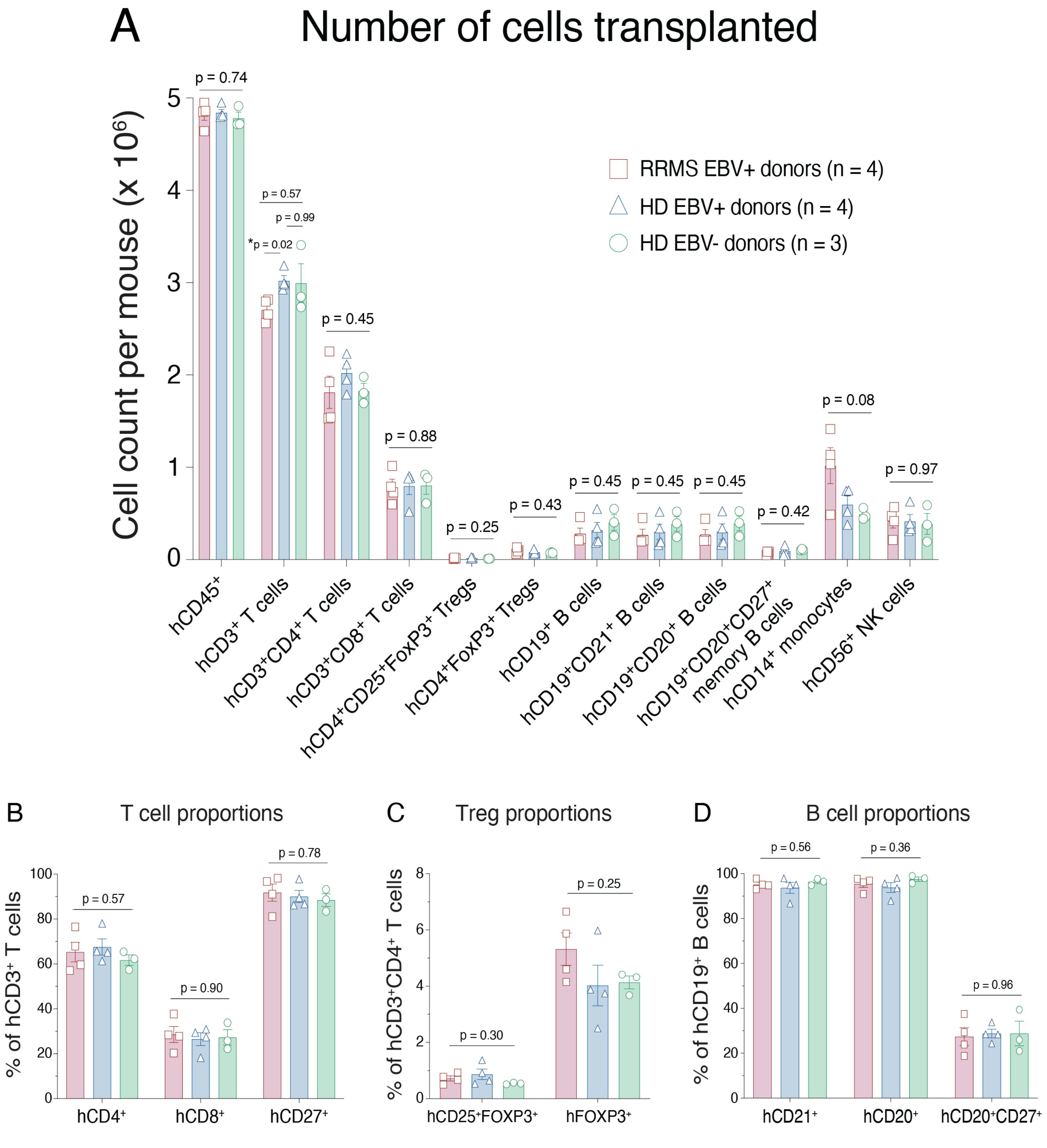

A
Number of cells transplanted
Cell count per mouse (x 10$^6$)
RRMS EBV+ donors (n = 4)
HD EBV+ donors (n = 4)
HD EBV- donors (n = 3)
p = 0.74
p = 0.57
p = 0.99
*p = 0.02
p = 0.45
p = 0.88
p = 0.25
p = 0.43
p = 0.45
p = 0.45
p = 0.45
p = 0.42
p = 0.08
p = 0.97
0
1
2
3
4
5
hCD45$^+$
hCD3$^+$ T cells
hCD3$^+$CD4$^+$ T cells
hCD3$^+$CD8$^+$ T cells
hCD4$^+$CD25$^+$FoxP3$^+$ Tregs
hCD4$^+$FoxP3$^+$ Tregs
hCD19$^+$ B cells
hCD19$^+$CD21$^+$ B cells
hCD19$^+$CD20$^+$ B cells
hCD19$^+$CD20$^+$CD27$^+$ memory B cells
hCD14$^+$ monocytes
hCD56$^+$ NK cells
B
T cell proportions
% of hCD3$^+$ T cells
p = 0.57
p = 0.90
p = 0.78
0
20
40
60
80
100
hCD4$^+$
hCD8$^+$
hCD27$^+$
C
Treg proportions
% of hCD3$^+$CD4$^+$ T cells
p = 0.30
p = 0.25
0
2
4
6
8
hCD25$^+$FOXP3$^+$
hFOXP3$^+$
D
B cell proportions
% of hCD19$^+$ B cells
p = 0.56
p = 0.36
p = 0.96
0
20
40
60
80
100
hCD21$^+$
hCD20$^+$
hCD20$^+$CD27$^+$

### Figure S3

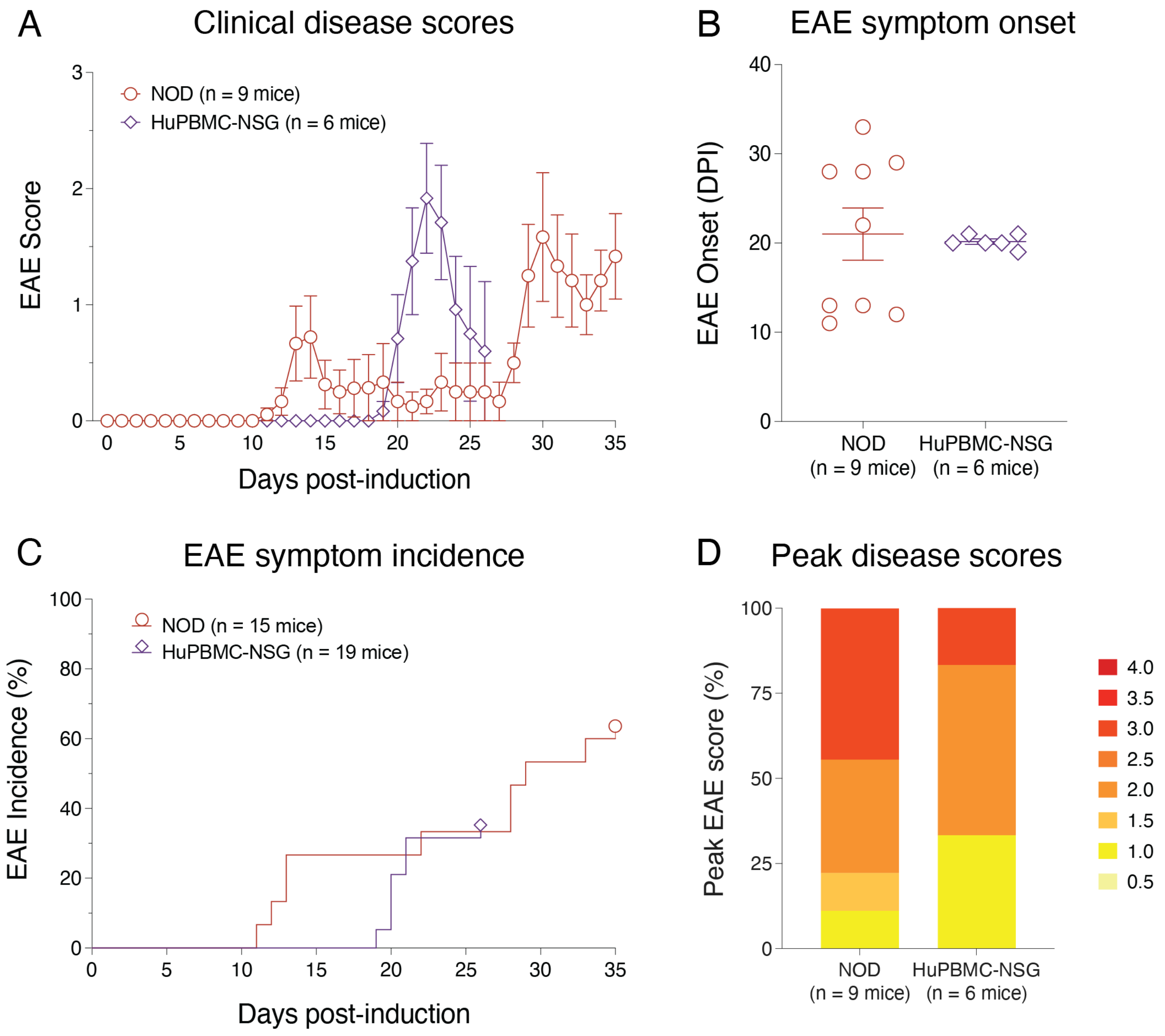
A
Clinical disease scores
NOD (n = 9 mice)
HuPBMC-NSG (n = 6 mice)
EAE Score
Days post-induction
B
EAE symptom onset
EAE Onset (DPI)
NOD
(n = 9 mice)
HuPBMC-NSG
(n = 6 mice)
C
EAE symptom incidence
NOD (n = 15 mice)
HuPBMC-NSG (n = 19 mice)
EAE Incidence (%)
Days post-induction
D
Peak disease scores
Peak EAE score (%)
NOD
(n = 9 mice)
HuPBMC-NSG
(n = 6 mice)
4.0
3.5
3.0
2.5
2.0
1.5
1.0
0.5

### Figure S4

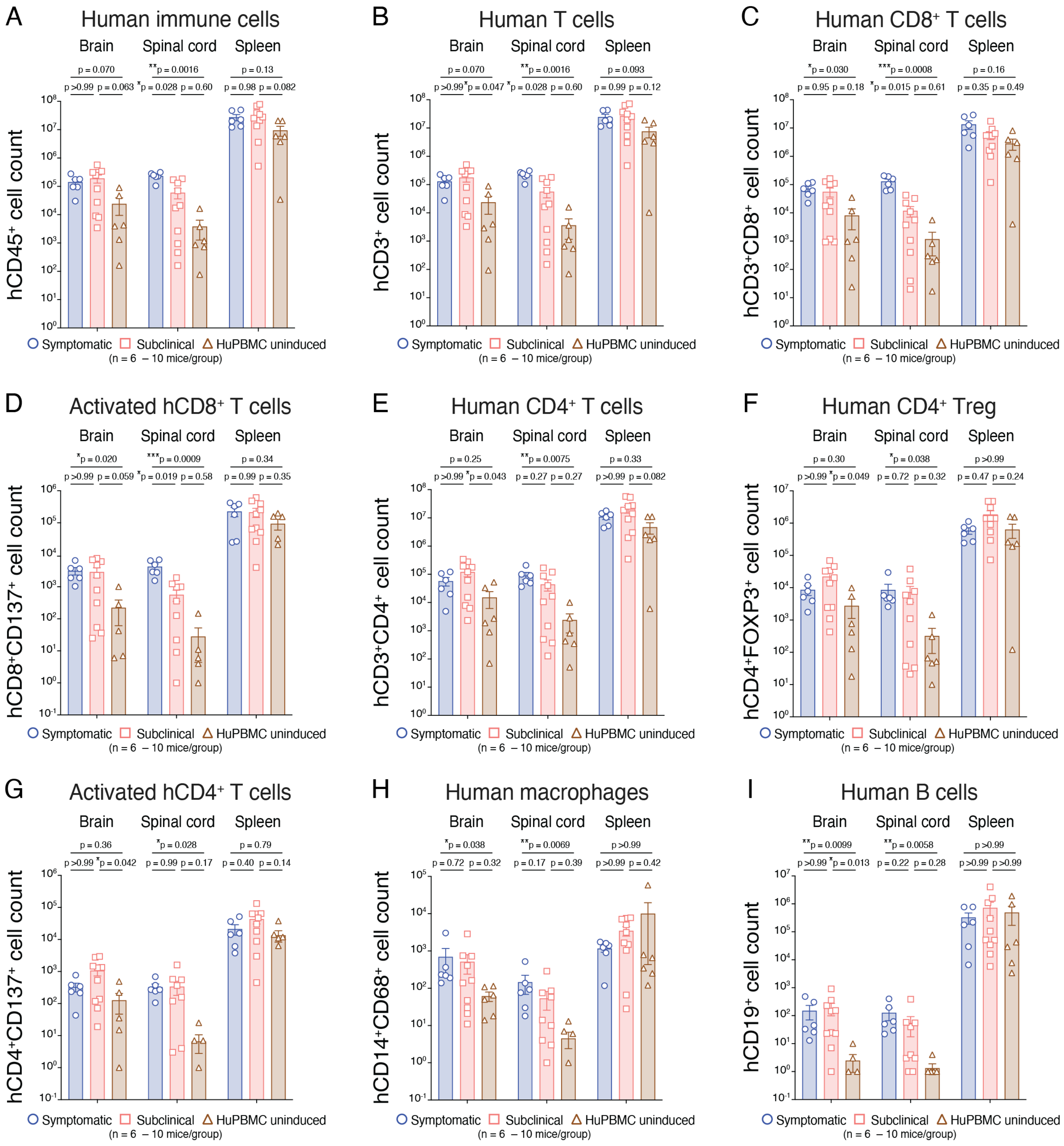
A
Human immune cells
Brain
Spinal cord
Spleen
p = 0.070
**p = 0.0016
p = 0.13
p >0.99
p = 0.063
*p = 0.028
p = 0.60
p = 0.98
p = 0.082
hCD45+ cell count
B
Human T cells
Brain
Spinal cord
Spleen
p = 0.070
**p = 0.0016
p = 0.093
p >0.99
*p = 0.047
*p = 0.028
p = 0.60
p = 0.99
p = 0.12
hCD3+ cell count
C
Human CD8+ T cells
Brain
Spinal cord
Spleen
*p = 0.030
***p = 0.0008
p = 0.16
p = 0.95
p = 0.18
*p = 0.015
p = 0.61
p = 0.35
p = 0.49
hCD3+CD8+ cell count
D
Activated hCD8+ T cells
Brain
Spinal cord
Spleen
*p = 0.020
***p = 0.0009
p = 0.34
p >0.99
p = 0.059
*p = 0.019
p = 0.58
p = 0.99
p = 0.35
hCD8+CD137+ cell count
E
Human CD4+ T cells
Brain
Spinal cord
Spleen
p = 0.25
**p = 0.0075
p = 0.33
p >0.99
*p = 0.043
p = 0.27
p = 0.27
p >0.99
p = 0.082
hCD3+CD4+ cell count
F
Human CD4+ Treg
Brain
Spinal cord
Spleen
p = 0.30
*p = 0.038
p >0.99
p >0.99
*p = 0.049
p = 0.72
p = 0.32
p = 0.47
p = 0.24
hCD4+FOXP3+ cell count
G
Activated hCD4+ T cells
Brain
Spinal cord
Spleen
p = 0.36
*p = 0.028
p = 0.79
p >0.99
*p = 0.042
p = 0.99
p = 0.17
p = 0.40
p = 0.14
hCD4+CD137+ cell count
H
Human macrophages
Brain
Spinal cord
Spleen
*p = 0.038
**p = 0.0069
p >0.99
p = 0.72
p = 0.32
p = 0.17
p = 0.39
p >0.99
p = 0.42
hCD14+CD68+ cell count
I
Human B cells
Brain
Spinal cord
Spleen
**p = 0.0099
**p = 0.0058
p >0.99
p >0.99
*p = 0.013
p = 0.22
p = 0.28
p >0.99
p >0.99
hCD19+ cell count
Symptomatic
Subclinical
HuPBMC uninduced
(n = 6 – 10 mice/group)

### Figure S5

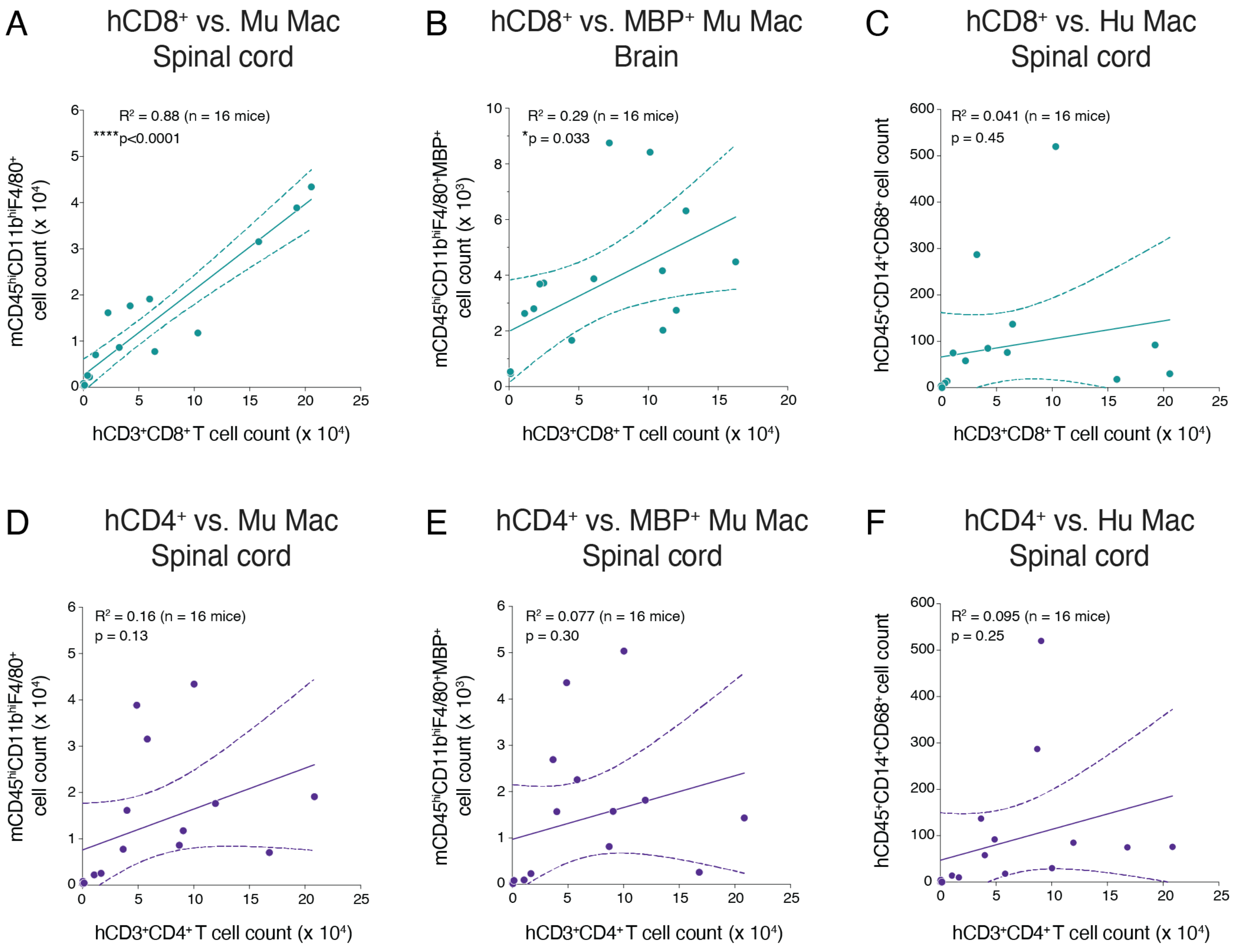
A
hCD8⁺ vs. Mu Mac
Spinal cord
R² = 0.88 (n = 16 mice)
****p<0.0001
mCD45ʰⁱCD11bʰⁱF4/80⁺ cell count (x 10⁴)
hCD3⁺CD8⁺ T cell count (x 10⁴)
B
hCD8⁺ vs. MBP⁺ Mu Mac
Brain
R² = 0.29 (n = 16 mice)
*p = 0.033
mCD45ʰⁱCD11bʰⁱF4/80⁺MBP⁺ cell count (x 10³)
hCD3⁺CD8⁺ T cell count (x 10⁴)
C
hCD8⁺ vs. Hu Mac
Spinal cord
R² = 0.041 (n = 16 mice)
p = 0.45
hCD45⁺CD14⁺CD68⁺ cell count
hCD3⁺CD8⁺ T cell count (x 10⁴)
D
hCD4⁺ vs. Mu Mac
Spinal cord
R² = 0.16 (n = 16 mice)
p = 0.13
mCD45ʰⁱCD11bʰⁱF4/80⁺ cell count (x 10⁴)
hCD3⁺CD4⁺ T cell count (x 10⁴)
E
hCD4⁺ vs. MBP⁺ Mu Mac
Spinal cord
R² = 0.077 (n = 16 mice)
p = 0.30
mCD45ʰⁱCD11bʰⁱF4/80⁺MBP⁺ cell count (x 10³)
hCD3⁺CD4⁺ T cell count (x 10⁴)
F
hCD4⁺ vs. Hu Mac
Spinal cord
R² = 0.095 (n = 16 mice)
p = 0.25
hCD45⁺CD14⁺CD68⁺ cell count
hCD3⁺CD4⁺ T cell count (x 10⁴)

### Figure S6

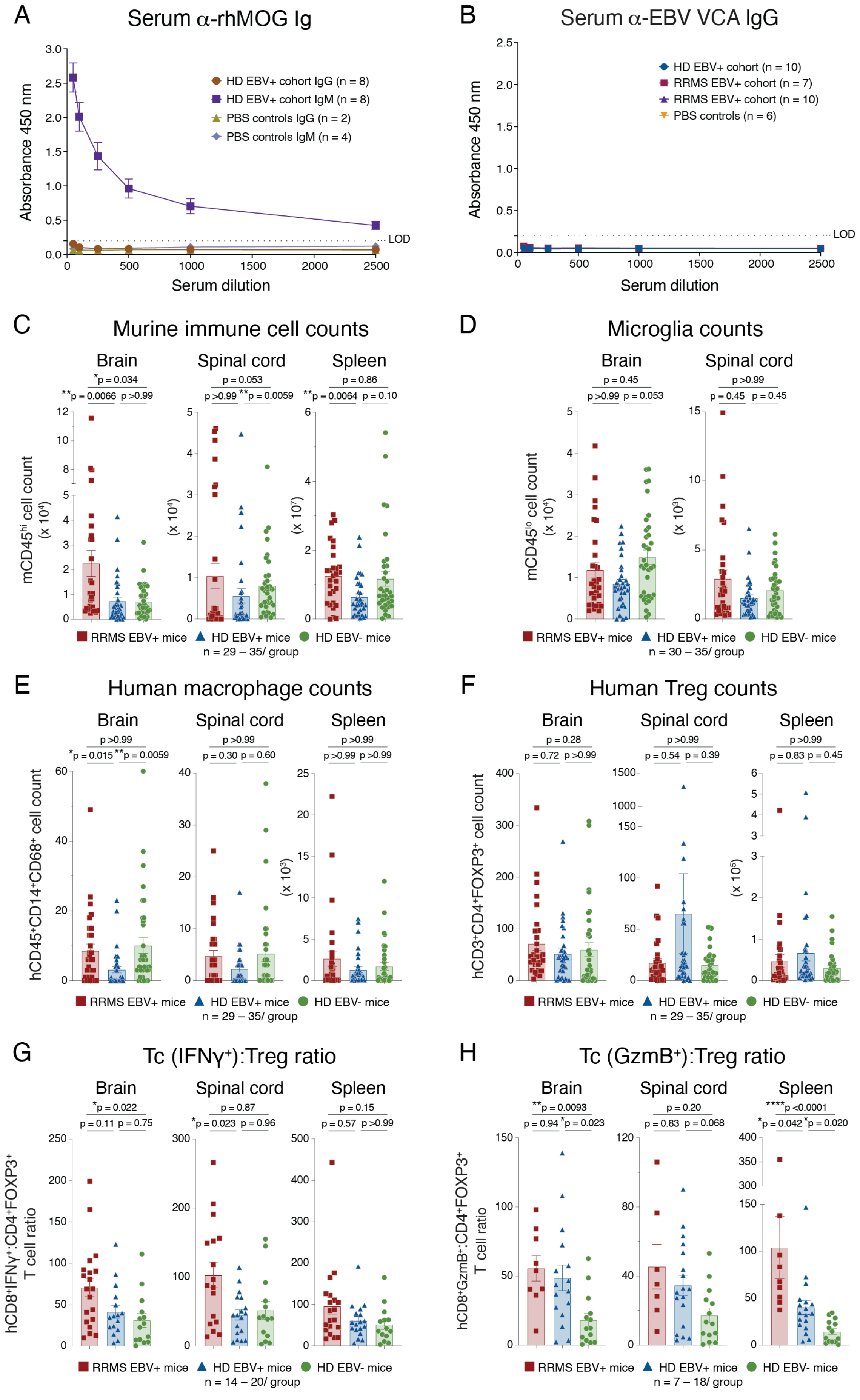

A
Serum α-rhMOG Ig
Absorbance 450 nm
Serum dilution
HD EBV+ cohort IgG (n = 8)
HD EBV+ cohort IgM (n = 8)
PBS controls IgG (n = 2)
PBS controls IgM (n = 4)
LOD
B
Serum α-EBV VCA IgG
HD EBV+ cohort (n = 10)
RRMS EBV+ cohort (n = 7)
RRMS EBV+ cohort (n = 10)
PBS controls (n = 6)
C
Murine immune cell counts
Brain
Spinal cord
Spleen
mCD45hi cell count
RRMS EBV+ mice
HD EBV+ mice
HD EBV- mice
n = 29 – 35/ group
D
Microglia counts
mCD45lo cell count
n = 30 – 35/ group
E
Human macrophage counts
hCD45+CD14+CD68+ cell count
n = 29 – 35/ group
F
Human Treg counts
hCD3+CD4+FOXP3+ cell count
n = 29 – 35/ group
G
Tc (IFNγ+):Treg ratio
hCD8+IFNγ+:CD4+FOXP3+ T cell ratio
n = 14 – 20/ group
H
Tc (GzmB+):Treg ratio
hCD8+GzmB+:CD4+FOXP3+ T cell ratio
n = 7 – 18/ group

Figure S7

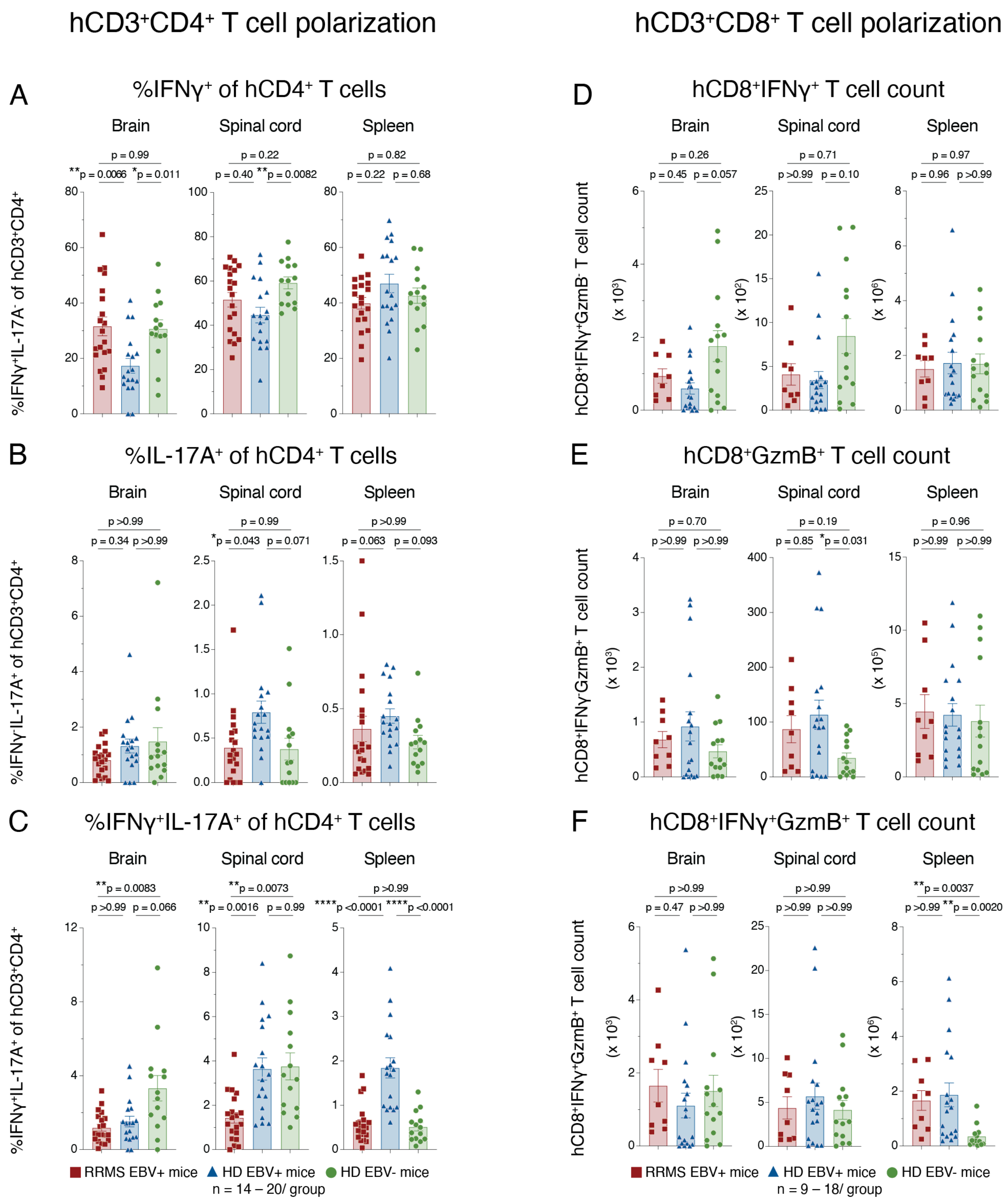
hCD3+CD4+ T cell polarization
hCD3+CD8+ T cell polarization
A
%IFNγ+ of hCD4+ T cells
Brain
Spinal cord
Spleen
p = 0.99
**p = 0.0066 *p = 0.011
p = 0.22
p = 0.40 **p = 0.0082
p = 0.82
p = 0.22 p = 0.68
%IFNγ+IL-17A- of hCD3+CD4+
B
%IL-17A+ of hCD4+ T cells
Brain
Spinal cord
Spleen
p >0.99
p = 0.34 p >0.99
p = 0.99
*p = 0.043 p = 0.071
p >0.99
p = 0.063 p = 0.093
%IFNγ-IL-17A+ of hCD3+CD4+
C
%IFNγ+IL-17A+ of hCD4+ T cells
Brain
Spinal cord
Spleen
**p = 0.0083
p >0.99 p = 0.066
**p = 0.0073
**p = 0.0016 p = 0.99
p >0.99
****p <0.0001 ****p <0.0001
%IFNγ+IL-17A+ of hCD3+CD4+
RRMS EBV+ mice
HD EBV+ mice
HD EBV- mice
n = 14 – 20/ group
D
hCD8+IFNγ+ T cell count
Brain
Spinal cord
Spleen
p = 0.26
p = 0.45 p = 0.057
p = 0.71
p >0.99 p = 0.10
p = 0.97
p = 0.96 p >0.99
hCD8+IFNγ+GzmB- T cell count
(x 10³)
(x 10²)
(x 10⁶)
E
hCD8+GzmB+ T cell count
Brain
Spinal cord
Spleen
p = 0.70
p >0.99 p >0.99
p = 0.19
p = 0.85 *p = 0.031
p = 0.96
p >0.99 p >0.99
hCD8+IFNγ-GzmB+ T cell count
(x 10³)
(x 10⁵)
F
hCD8+IFNγ+GzmB+ T cell count
Brain
Spinal cord
Spleen
p >0.99
p = 0.47 p >0.99
p >0.99
p >0.99 p >0.99
**p = 0.0037
p >0.99 **p = 0.0020
hCD8+IFNγ+GzmB+ T cell count
(x 10³)
(x 10²)
(x 10⁶)
RRMS EBV+ mice
HD EBV+ mice
HD EBV- mice
n = 9 – 18/ group

### Figure S8

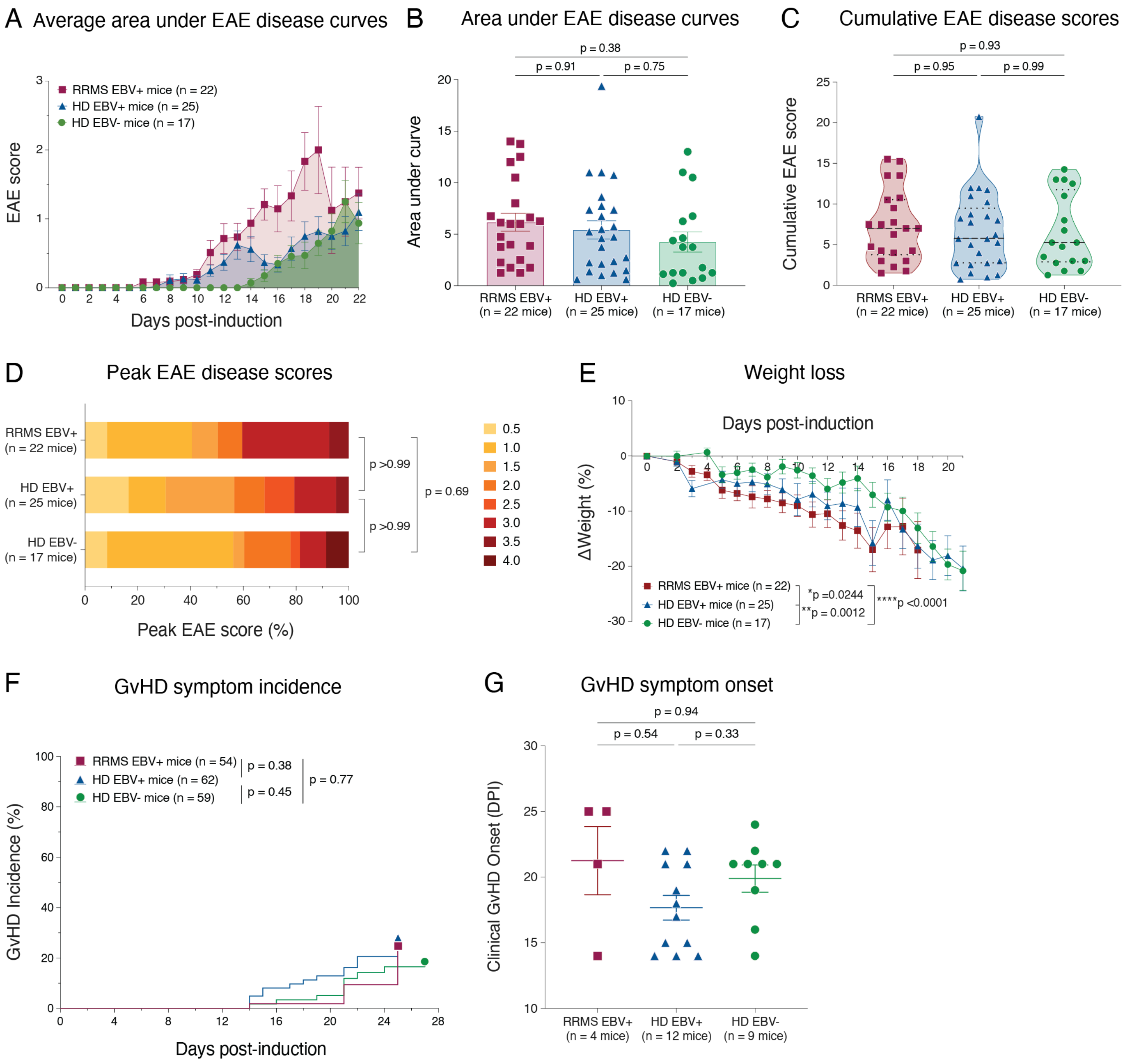
A Average area under EAE disease curves
RRMS EBV+ mice (n = 22)
HD EBV+ mice (n = 25)
HD EBV- mice (n = 17)
EAE score
Days post-induction
B Area under EAE disease curves
p = 0.38
p = 0.91
p = 0.75
Area under curve
RRMS EBV+ (n = 22 mice)
HD EBV+ (n = 25 mice)
HD EBV- (n = 17 mice)
C Cumulative EAE disease scores
p = 0.93
p = 0.95
p = 0.99
Cumulative EAE score
D Peak EAE disease scores
RRMS EBV+ (n = 22 mice)
HD EBV+ (n = 25 mice)
HD EBV- (n = 17 mice)
p >0.99
p = 0.69
p >0.99
Peak EAE score (%)
0.5
1.0
1.5
2.0
2.5
3.0
3.5
4.0
E Weight loss
Days post-induction
ΔWeight (%)
RRMS EBV+ mice (n = 22)
HD EBV+ mice (n = 25)
HD EBV- mice (n = 17)
*p =0.0244
**p = 0.0012
****p <0.0001
F GvHD symptom incidence
RRMS EBV+ mice (n = 54)
HD EBV+ mice (n = 62)
HD EBV- mice (n = 59)
p = 0.38
p = 0.45
p = 0.77
GvHD Incidence (%)
Days post-induction
G GvHD symptom onset
p = 0.94
p = 0.54
p = 0.33
Clinical GvHD Onset (DPI)
RRMS EBV+ (n = 4 mice)
HD EBV+ (n = 12 mice)
HD EBV- (n = 9 mice)

### Figure S9

#### Donor PBMC T cell compositiion after freeze-thaw

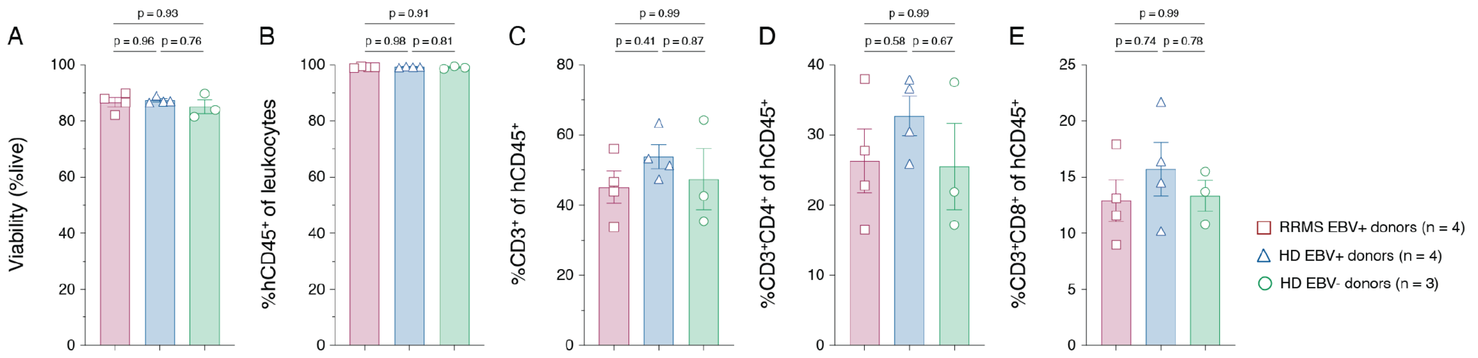


#### Baseline donor T cell activation marker expression (untreated PBMC)

##### hCD3⁺CD4⁺ T cells

##### hCD3⁺CD8⁺ T cells

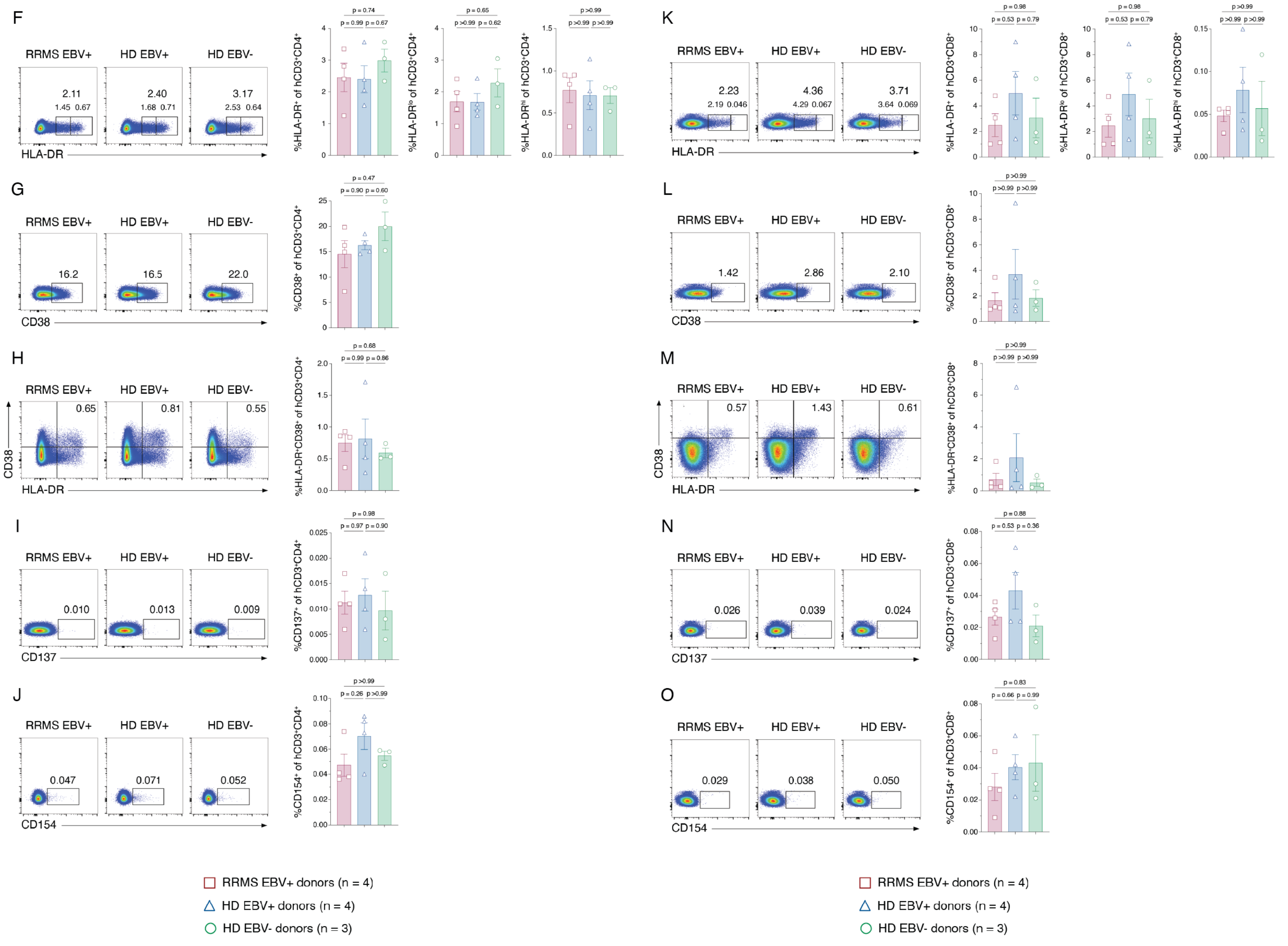

### Figure S10

#### Basline donor T cell polarization

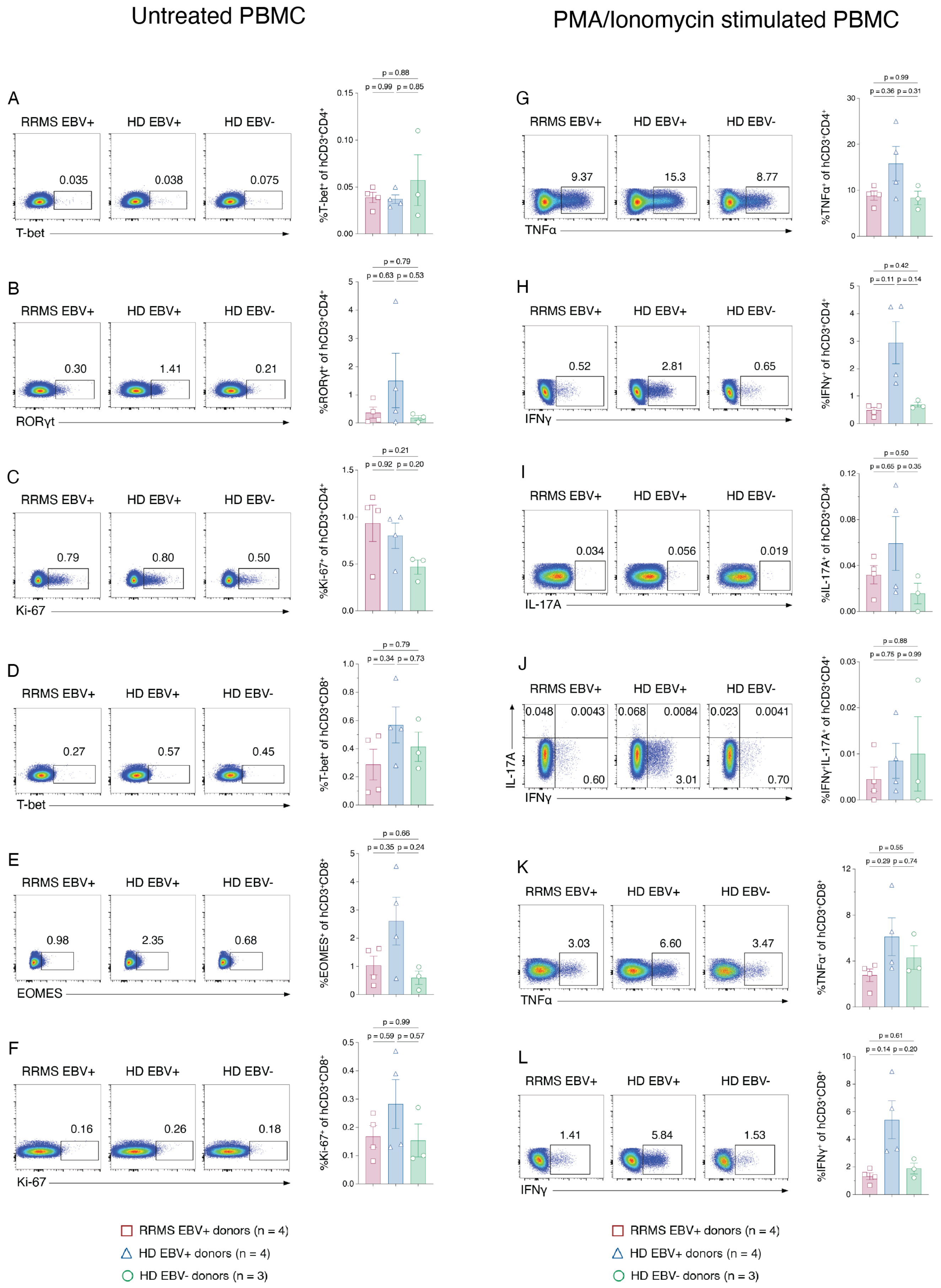
